# Divergent mechanics of inner ear morphogenesis are coupled to developmental tempo over vertebrate evolution

**DOI:** 10.64898/2026.02.27.706697

**Authors:** Shunya Kuroda, Shuhei A. Horiguchi, Soichiro Kato, Oki Hayasaka, Hiroyasu Kamei, Wataru Takagi, Shinnosuke Higuchi, Tatsuya Hirasawa, Koji Fujimura, Kiyoshi Hiraoka, Nanoka Suzuki, Makoto Suzuki, Hajime Ogino, Anna Yasunaga, Hiroshi Kiyonari, Shigeru Kuratani, Satoru Okuda

## Abstract

Vertebrate embryos vary in size and developmental tempo based on reproductive strategies; however, organs of ancient origin, such as the eye and inner ear, remain highly conserved in structure and function. How both diverse embryonic traits and conserved organ development occur remains unclear. Here, we identify divergent morphogenetic mechanics of inner ear development, particularly in vesicular primordium expansion, in 12 species representing the major vertebrate lineages. Differences in cell volume regulation determine tissue growth mechanics, which diversify the expansion process but yield a conserved morphological outcome. Heterochronic shifts in organogenesis allow this regulation to evolve in coordination with a prolonged developmental period. These findings suggest that alternative morphogenetic mechanics mitigate the effects of diversified embryonic traits, thereby ensuring consistent morphological outcomes for highly conserved organs.

## Main Text

Vertebrates have undergone remarkable diversification in forms, sizes, and reproductive modes to adapt to diverse environments, ranging from aquatic and terrestrial habitats to parasitic lifestyles (*1, 2*). This adaptation was accompanied by the evolution of varied developmental strategies (*3*), which may have necessitated modifications of ancestral morphogenetic processes. For example, the evolution of large and yolky eggs, which arose in response to selective factors such as high predation (*1*), mechanically constrains egg cleavage and results in divergent modes of early development (*4, 5*). Likewise, evolutionary changes in developmental timing, such as paedomorphosis in salamanders that inhabit environments where remaining aquatic is preferable to transitioning to land (*6*), can modify morphogenesis. For instance, these changes may cause the loss of facial bones, such as the maxilla in *Necturus* (*7*). This view, that morphological change arises from shifts in the relative timing of developmental events, is rooted in the classical concept of heterochrony originated by Haeckel [(*8*), reviewed by ref (*3*)].

In contrast to these extensive evolutionary modifications, the fundamental components of the vertebrate body plan—including the eye, inner ear, and tubular nerve cord—have remained highly conserved throughout evolution (*9, 10*). This raises the question of whether the morphogenetic processes of these organs may have endured conflict between the diversification of developmental strategies and the preservation of morphological outcomes. Such a conflict has been proposed to give rise to developmental system drift (DSD) (*11*), in which the evolutionary modifications of developmental processes occur while maintaining homologous outcomes, under conflicting directional and stabilizing selection pressures (*12, 13*). We hypothesize that DSD mitigates the effects of diversified embryonic traits, such as embryo size and developmental timing, thereby stabilizing morphological outcomes in conserved organs. Given this hypothesis, a central question is how vertebrates have acquired various developmental strategies while retaining conserved organ morphology through DSD.

To approach this question, we focused on the inner ear, an organ that has remained conserved despite the diversification of vertebrate developmental strategies. The inner ear is a balance-sensing organ present in all extant and presumably fossil vertebrates (*14*) and contains semicircular canals that detect rotational head movements (*15*). Although the number of canals varies among species, each canal retains a conserved toroidal geometry and mechanosensory function (*15-17*). The conserved geometry and function presumably reflect long-standing phenotypic stabilizing selection (*3, 18*) occurring over more than 500 million years of vertebrate evolution. During this period, embryonic traits have diversified dramatically, with egg diameters ranging from approximately 100 μm in mammals to greater than 10 cm in ostriches (*19, 20*) and developmental durations spanning from days in zebrafish to years in deep-sea sharks (*21, 22*). Therefore, the inner ear is an ideal system for investigating how embryos with diverse developmental strategies reliably generate a conserved structure through development and evolution.

Describing DSD inevitably involves a mechanistic understanding of how divergent developmental processes yield a conserved outcome (*12*). Although evolutionary changes in developmental processes occur at multiple levels—from morphological to genetic—here, we focus on morphogenetic mechanics (*23, 24*), the physical processes that can evolve in response to various developmental strategies and are the immediate control of tissue geometry (*25*). From a mechanical perspective, inner ear morphology arises through a stereotyped morphogenetic sequence in which the otic vesicle (OTV) first expands and then forms protrusions that fuse to form pillars of semicircular canals (*26*). In zebrafish, OTV expansion is driven by hydrostatic pressure from the lumen and viscoelastic deformation of the epithelial wall (*27*). However, it remains unknown whether this mechanical sequence of inner ear morphogenesis—well characterized in zebrafish—is conserved across vertebrates, including non-model animals.

Here, we investigate how the mechanics of inner ear morphogenesis vary across vertebrates with divergent developmental strategies. By linking interspecies differences in morphogenetic mechanics to developmental strategies, we aim to uncover how developmental systems reconcile broad diversification with conserved morphogenetic outcomes across development and evolution.

## Results

### Two modes of inner ear morphogenesis across vertebrates

To identify how morphogenetic mechanics diversify across vertebrates, we first compared the overall morphogenetic sequence of inner ear formation among species. Inner ear morphogenesis proceeds in two sequential phases: an initial OTV expansion followed by semicircular canal formation (fig. S1). During canal formation, the OTV epithelium develops protrusions with shapes that vary depending on species— finger-like protrusions in zebrafish and frogs and bowl-like protrusions in mice and chickens (*28*).

To classify the protrusion shape across vertebrates, we parameterized the shapes by their width-to-depth ratio ω (fig. S2A) and measured it in 12 species representing all major vertebrate classes (as a taxonomic rank) (Fig. 1A, fig. S1). This analysis revealed that the protrusion shape tended toward either a finger-like or a bowl-like shapes (Fig. 1B, C, fig. S2B). We next examined diversity in the OTV expansion phase, which has not been comprehensively described before. We parameterized vesicle shape by epithelial thickness and identified two broad trends (Fig. 1D, figs. S1 and S3): in one, the epithelium thinned significantly, whereas in the other, it thickened (Fig. 1E). Notably, the mode of OTV expansion showed a general correlation with that of subsequent canal formation (Fig. 1F). Specifically, species that undergo epithelial thinning during OTV expansion tend to have finger-like protrusions, whereas species that undergo epithelial thickening, or at least do not demonstrate thinning during OTV expansion, are more likely to form bowl-like protrusions.

**Fig. 1.**
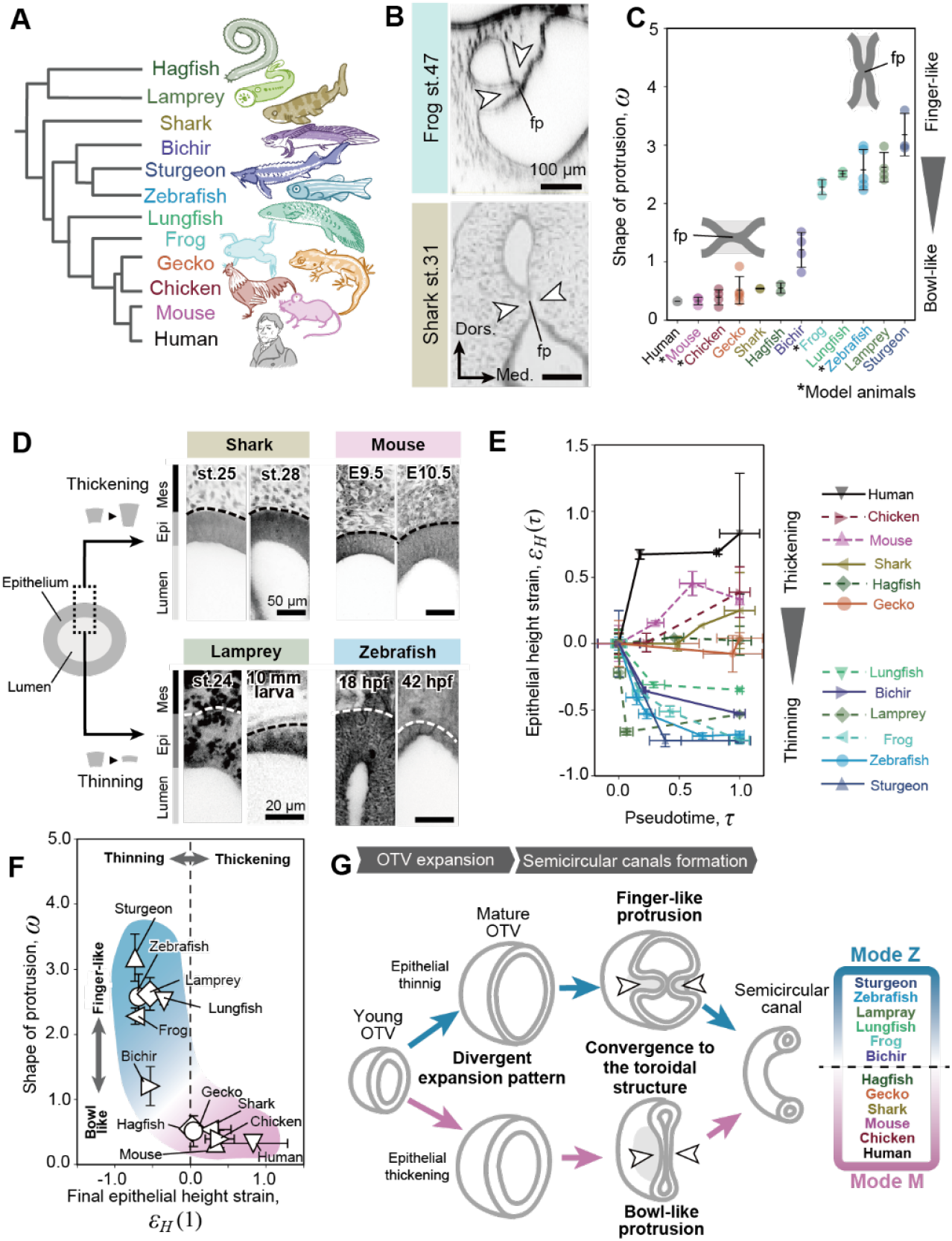
Two discrete morphogenetic processes in inner ear development. (**A**) Schematic tree showing the phylogenetic relationships of animals investigated in this study. (**B**) Representative confocal (top, inverted) and hematoxylin and eosin (bottom, greyscale) sectional images showing paired protrusion of OTV wall (arrowheads) contacts as a fusion plate in a frog and a shark embryo. (**C**) Quantification of the shape of protrusions *ω* among species (*N* = 1–5 depending on species and stages, see table S1). (**D**) Examples of wall thickness change from young to mature OTVs in representative species. Embryos were stained with RBITC (inverted). Dashed lines delineate the basal surface of the OTV. (**E**) Temporal changes in the epithelial height over pseudotime *τ* (*N* = 1–6 depending on species and stages, see fig. S1). (**F**) Data from C and E plotted in a 2D parameter space to show diversity in morphogenesis of the vertebrate inner ear. (**G**) Schematic representation of two modes (Z and M) of morphogenetic processes found in the inner ear. Arrowheads show a pair of protrusions. The younger stage samples of hagfish and lungfish may be partially incomplete because of limited sample accessibility. Each datapoint in C, E, and F represents the mean of one of the 12 species collected; error bars indicate means ± SDs. Dors., dorsal; fp, fusion plate; Med., medial. Scale bars = 100 µm (C), 50 µm (E top), and 20 µm (E bottom).

These results indicate that inner ear morphogenesis follows divergent processes spanning two extreme modes (hereafter referred to as Z- and M-modes, named after zebrafish and mouse, respectively), revealing previously unrecognized DSD in inner ear formation (Fig. 1G). For subsequent analyses, we classified the 12 examined species into those exhibiting either the Z- or M-mode primarily based on the temporal changes in epithelial thickness: species with a final epithelial height strain *ε*_*H*_(1) < 0 were classified as Z-mode species, whereas those with *ε*_*H*_(1) > 0 were classified as M-mode species (Fig. 1F, G).

### Distinct mechanics of OTV expansion underlying the two morphogenetic modes

To investigate the mechanical basis of divergent OTV expansion, we used zebrafish and mice as experimental model organisms of the Z- and M-mode species, respectively, and characterized their morphogenetic processes in terms of geometry and mechanics. From a geometrical perspective, OTV expansion can be described as a combination of (i) an increase in lumen volume (*V*_*L*_) due to fluid accumulation and (ii) an increase in epithelial tissue volume (*V*_*E*_) driven by cell proliferation and/or hypertrophy (Fig. 2A). From a mechanical perspective, vesicle geometry is maintained by a balance of forces at each time point: luminal pressure (*P*) resisting compression of the luminal fluid and tensile stress (σ) resisting in-plane stretching of the epithelium (Fig. 2B).

**Fig. 2.**
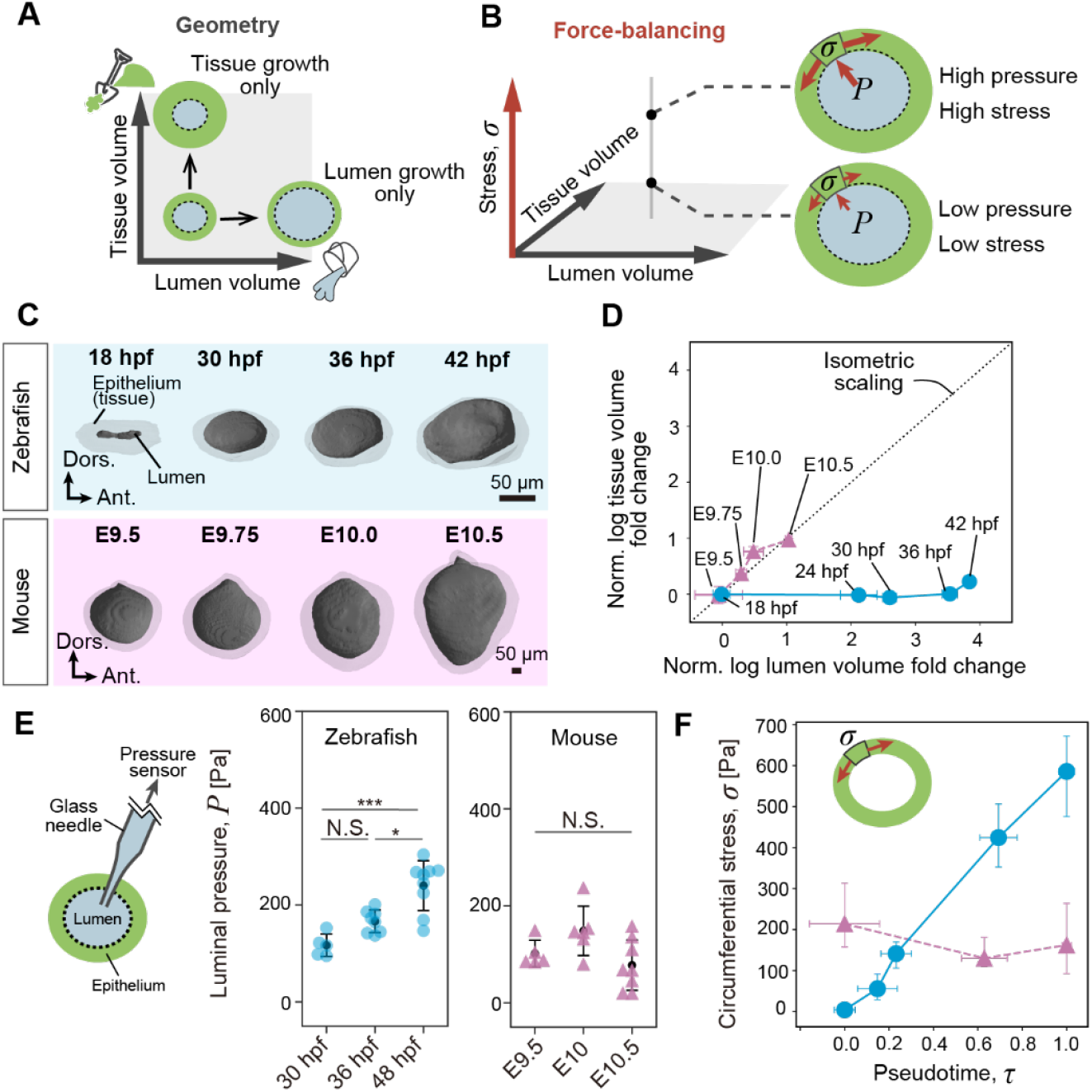
Two discrete OTV growth mechanics in zebrafish and mice. (**A, B**) Diagrams showing the concept of geometrical (A) and mechanical (B) documentations of the OTVs. (**C**) 3D reconstructions of OTVs along expansion processes in zebrafish (18–42 hpf) and mice (E9.5–10.5). (**D**) Temporal changes in OTV geometry in zebrafish (solid lines; *N* = 4–6) and mice (dashed lines; *N* = 2–9), expressed as the normalized tissue versus lumen volume fold change. The dotted line (*y* = *x*) represents ideal isometric scaling. (**E**) Measurement of luminal pressure in zebrafish [circles; data are from ref.(*27*))] and mice (triangles) at each stage (zebrafish; *N* = 5, 8, 9, mice; *N* = 5, 6, 8). Turkey’s test (zebrafish), Kruskal-Wallis test (mice). \**P* < 0.05, \*\*\**P* < 0.001, N.S. = not significant. (**F**) Circumferential stress as a function of pseudotime in zebrafish (solid line) and mice (dashed line). Error bars in D and E are ± SDs. Error bars of circumferential stress in F represents 95% confidence intervals. Ant., anterior; Dors., dorsal. Scale bars = 50 µm (C).

First, we quantified temporal changes in the epithelial tissue volume in zebrafish (18-42 hours post fertilization, hpf) and mice (embryonic day [E]9.5–10.5). Tissue volumetric growth was limited in zebrafish, particularly until 36 hpf, whereas continuous tissue growth occurred in mice (Fig. 2C, D). Next, we quantified the tissue-to-lumen volume ratio, which reflects overall OTV geometry during expansion. Intriguingly, this ratio remained constant in mice but decreased in zebrafish (Fig. 2D). These results indicate that OTV expansion in mice proceeds with isometric scaling (*V*_*E*_ ∝ *V*_*L*_) (*29*) accompanied by epithelial thickening, whereas in zebrafish, expansion is driven mainly by luminal enlargement with little change in tissue volume, resulting in epithelial thinning.

From a mechanical perspective, a pioneering study by Mosaliganti and colleagues using zebrafish (*27*) demonstrated that luminal pressure increases during OTV expansion. To test whether the mouse OTV displays the same trend, we measured luminal pressure using a custom-built device (fig. S4A, B). In contrast to zebrafish, luminal pressure in mice remained nearly constant at ∼100 Pa throughout OTV expansion (Fig. 2E, fig. S4C). Using the Young–Laplace equation (*30*), we estimated the circumferential tensile stress in the epithelium and found that the stress in the mouse OTV is maintained within a narrow range (∼200 Pa), in contrast to the progressively increasing stress observed in zebrafish (Fig. 2F).

These results highlight distinct mechanical processes underlying OTV expansion: in zebrafish, expansion is dominated by luminal inflation, with luminal pressure and epithelial stress increasing in parallel; in mice, expansion involves concurrent enlargement of both the lumen and tissue volumes while maintaining constant luminal pressure and epithelial stress.

### Mechanosensitive epithelial growth diversifies morphogenetic mechanics

Motivated by our observation that mouse OTVs expand while maintaining nearly constant epithelial stress (Fig. 2F), we hypothesized that this divergence arises from differences in mechanosensitive cellular responses (*31*). Specifically, an active, mechanosensitive epithelial growth mechanism might account for stress homeostasis, as previously reported for chicken neural tube expansion (*32*). We made two assumptions regarding this hypothesis: (i) luminal dynamics operates similarly in both species in that fluid transport into the lumen increases luminal volume, stretching the epithelium and thereby generating tensile stress and elevating luminal pressure, and (ii) the tissue response, in contrast, is species-specific in that in zebrafish, the epithelium remains elastically stretched under this increased tension, whereas in mice, cells respond to the increased tensile stress by promoting their volumetric growth, thereby relaxing the stress.

On the basis of this hypothesis, we constructed a mathematical model (Fig. 3A; see also Materials and Methods for details). In this model, tissue growth is regulated by a stress sensitivity coefficient *β*, which quantifies the mechanosensitive responses in epithelia. This model quantitatively recapitulated the observed temporal changes in *V*_*L*_, *V*_*E*_, and σ in both species (Fig 3B). Model fitting revealed that in zebrafish, *β* ≈ 0, indicating arrested epithelial tissue growth, whereas in mice, *β* > 0, which enables tissue growth to ensure stress homeostasis and isometric scaling. These analyses identified the stress sensitivity parameter *β* as the primary determinant for the divergence of OTV expansion modes, suggesting that mechanosensitive growth ability is a key factor that explains species-specific morphogenetic mechanics.

**Fig. 3.**
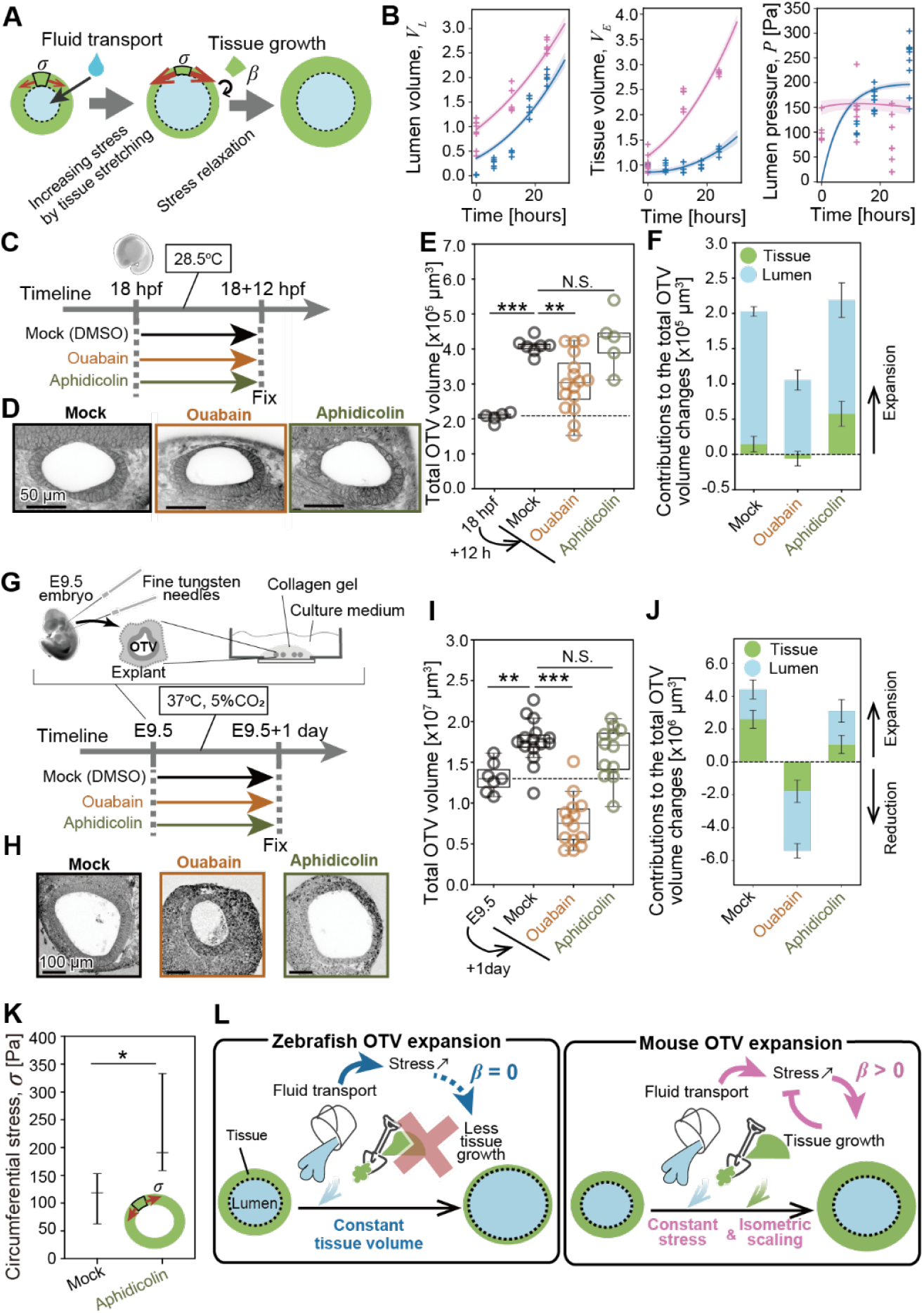
Pharmacological interventions in OTV expansion. (**A**) Diagram showing the concept of our mathematical model of OTV expansion. (**B**) Temporal changes in lumen volume (left), tissue volume (middle), and luminal pressure (right). Time on the horizontal axis represents hours after 18 hpf (zebrafish) or E9.5 (mice). Each plot represents measurements from fixed specimens, and lines show the results of fitting for zebrafish (blue) and mice (pink). (**C**) Time window of drug treatments in zebrafish. (**D**) Representative confocal images of zebrafish OTVs at 30 hpf treated only with DMSO (top), ouabain (middle), or aphidicolin (bottom). (**E**) Measured total OTV volume in embryos at 18 hpf, mock, and those treated with ouabain or aphidicolin (*N* = 5, 7, 16, 5). Games-Howell test. \*\**P* < 0.01, \*\*\**P* < 0.001, N.S., not significant. (**F**) Contributions of tissue (green) and lumen (blue) growth to the observed total OTV volume change for each experimental condition in zebrafish. (**G**) Schematic drawing of procedures for explant preparation and time window of drug treatments. (**H**) Representative confocal images of mouse OTV explants one day after treatment with DMSO (top), ouabain (middle), or aphidicolin (bottom). (**I**) Measured total OTV volume in embryos at E9.5, mock explants, and explants treated with ouabain or aphidicolin (*N* = 7, 15, 14, 12). Games-Howell test. \**P* < 0.05, \*\**P* < 0.01, \*\*\**P* < 0.001, N.S., not significant. (**J**) Contributions of tissue (green) and lumen (blue) growth to the total OTV volume change in each experimental condition in mouse explants are shown as stacked bar graphs. (**K**) Calculated circumferential stress of the mouse OTV explants after mock and aphidicolin treatment. Bootstrap hypothesis test for circumferential stress. \**P* < 0.05. (**L**) Schematic drawings showing species-specific expansion programs underlying two distinct OTV expansion processes in zebrafish and mice. Embryos and explants in D and H were stained with RBITC (inverted). In boxplots in E and I, the median (center horizontal line), range of data except for outliers (horizontal line), and range of the first to third quartiles (box) are indicated. Error bars in F and J indicate the mean ± SEs, and in K indicate 95% confidence intervals. Scale bars = 50 µm (D), 100 µm (H).

To verify the model predictions, we inhibited either fluid transport with ouabain (Na+/K+-pump inhibitor) or epithelial tissue growth with aphidicolin (by inhibiting the S-phase in the cell cycle) (*27, 33*). Zebrafish embryos at 18 hpf were treated for 12 hours with either of the inhibitors or the solvent alone (mock) (Fig. 3C). Ouabain treatment inhibited OTV expansion, whereas aphidicolin treatment had little effect on overall OTV size (Fig. 3D, E). The epithelial tissue volume remained unchanged by both treatments (fig. S5A). Under all conditions, luminal growth accounted for most of the total OTV expansion (Fig. 3F, fig. S5A–C). Thus, zebrafish OTV expansion is driven primarily by fluid transport, not epithelial growth.

For inhibitor experiments in mice, we developed a one-day OTV explant culture system using E9.5 embryos (Fig. 3G), which recapitulated *in vivo* OTV expansion involving both tissue and lumen growth (Fig. 3H–J, Movie S1). Ouabain treatment markedly reduced both luminal and epithelial tissue volumes (Fig. 3H–J, fig. S5D, E), supporting our hypothesis of mechanosensitive regulation of epithelial tissue growth in that reduced lumen growth decreases tensile stress, thereby decreasing tissue volumetric growth. In contrast, aphidicolin-treated explants achieved OTV expansion comparable to that in the mock condition (Fig. 3H–J, fig. S5D, F). In this treatment, luminal volume increased to the same level as that in the mock condition, indicating that fluid transport proceeds independently of epithelial tissue growth (fig. S5E). However, tissue growth was significantly suppressed (fig. S5D), and luminal pressure and tissue stress were elevated (Fig. 3K, fig. S5G–I), indicating that stress-dependent epithelial tissue growth is responsible for stress homeostasis.

These results reveal distinct mechanical regulatory pathways in zebrafish and mice, both captured by a shared mechanical framework for OTV expansion (Fig. 3L). The key distinction between the two species lies in their species-specific mechanosensitive responsiveness (*β*): a near-zero *β* in zebrafish leads to elastic deformation with constant epithelial tissue volume and increased stress, whereas a high *β* in mice results in elastoplastic deformation that enables stress relaxation, ensuring stress homeostasis and isometric scaling.

### Cell volume regulation underlies mechanosensitive epithelial growth

As the temporal changes of OTV geometry provide a clear signature for the distinct expansion mechanisms in zebrafish and mice (Figs. 2D and 3L), we analyzed these temporal changes in the remaining 10 species. This analysis revealed two distinct patterns, consistent with the classification of Z- and M-mode inner ear morphogenesis (Fig. 4A). Furthermore, by tuning species-specific parameters, our mathematical model successfully recapitulated the temporal changes in all examined species (Fig. 4B, fig. S6), suggesting that the divergent modes of OTV expansion seen among the examined vertebrate classes can be explained by a shared mechanical framework. In particular, estimated values of the stress-sensitive tissue growth parameter *β* directly account for the distinct modes of OTV expansion (Fig. 4C). Thus, the divergent morphogenetic modes classified based on the degree of epithelial thickening (Fig. 1E, G can be expressed in the diverse values of *β*.

**Fig. 4.**
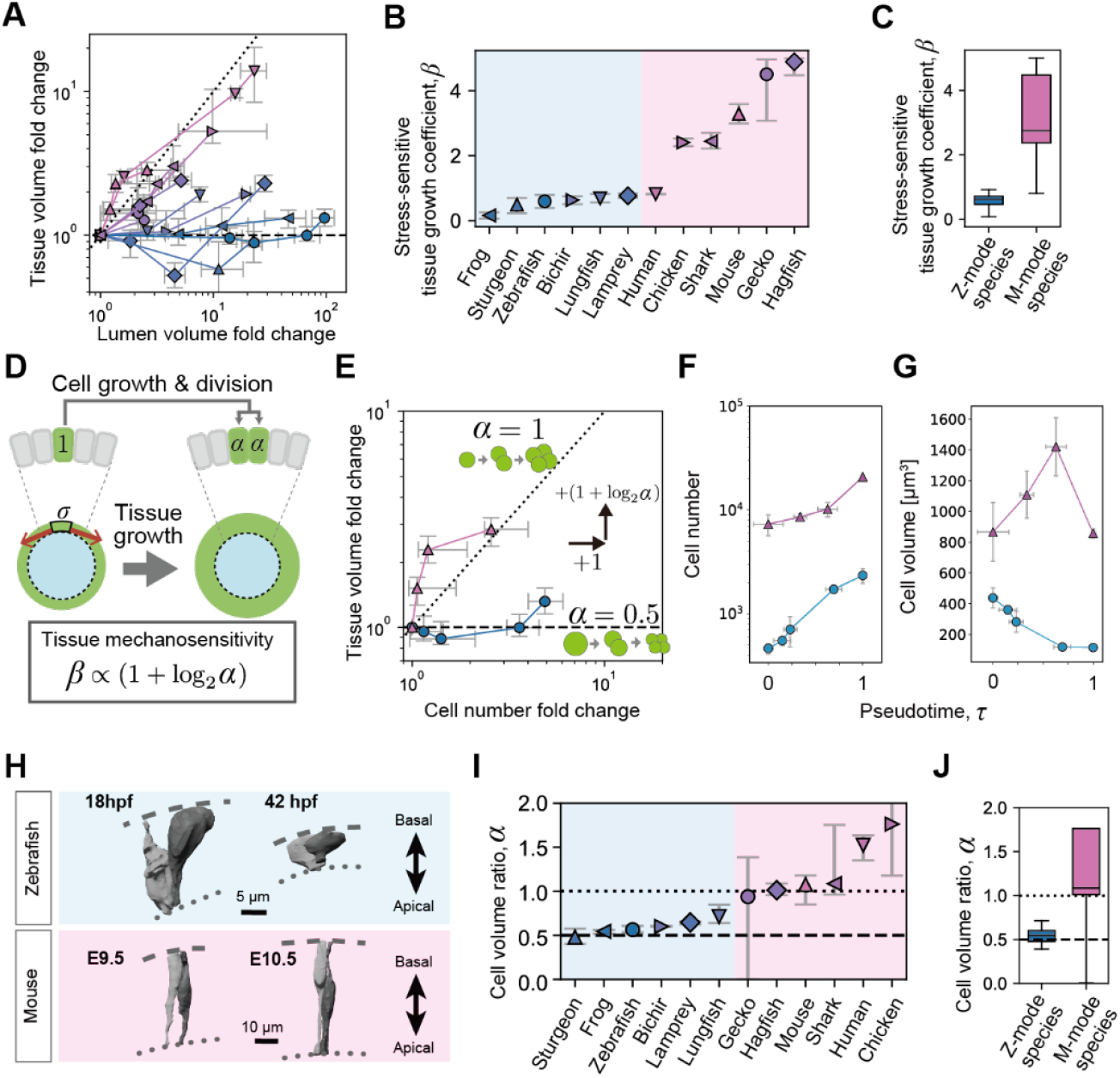
Differences in cell growth explain species-specific tissue growth patterns. (**A**) Cross-species comparison of temporal changes in OTV geometry plotted as lumen versus tissue volumes fold change. Z- and M-mode species tend to follow the constant-tissue-volume line (dashed) and the isometric scaling line (dotted), respectively. (**B, C**) Estimated tissue growth rate coefficient β by species (B) and aggregated by morphogenetic mode (C). (**D**) Schematic defining the intergenerational cell volume ratio α and linking cell growth to tissue growth. (**E**) Tissue volume as a function of cell number for zebrafish (blue) and mice (pink) on a log-log scale. The theory predicts that the slope equals 1 + log_2_ α (Materials and Methods). Zebrafish data approximately follow the dashed line (α = 0.5), whereas mouse data follow the dotted line (α = 1). (**F, G**) Changes in cell number (F) and mean cell volume (G) over developmental pseudotime τ (zebrafish; *N* = 4–6, mice; *N* = 2–9, depending on stage). (**H**) Representative 3D reconstructions of constituent cells of OTVs in zebrafish and mice. Dashed and dotted lines delineate basal and apical surfaces, respectively. (**I, J**) Estimated cell volume ratio α by species (I) and aggregated by mode of morphogenesis (J). Dashed and dotted lines denote α = 0.5 and α = 1, respectively. Points with error bars indicate the medians and 95% confidence intervals, except in F and G, which show means and SDs.

To elucidate the cellular basis of tissue growth responsiveness (*β*) during the OTV expansion period, we examined its relationship with cell volume and cell number. We quantified the contribution of cell volume regulation to tissue growth by introducing the intergenerational ratio of cell volume (*α*), defined as the mean daughter-to-mother ratio of cell volumes (Fig. 4D). We constructed a mathematical model describing the dynamics of cell volume and cell number and found that the epithelial tissue growth rate 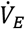, as well as *β*, is intrinsically scaled by 1 + log_2_ α (Fig. 4D; Materials and Methods). When α ≈ 1, the cell volume is maintained across generations, and tissue volume increases proportionally with cell number. In contrast, when α ≈ 0.5, cell divisions occur without net cell growth, resulting in constant tissue volume regardless of changes in the cell number (Fig. 4E). Thus, irrespective of the mechanosensitivity of cell number regulation, the mode of cell volume regulation largely governs the tissue-scale responsiveness to mechanical stimuli.

To test the theoretical prediction, we quantified the temporal changes in cell volume and cell number in representative species. Although tissue volumetric growth was limited in zebrafish (Fig. 2D), continuous cell division occurred in both zebrafish and mice (Fig. 4F). Moreover, cell volumes decreased in zebrafish but remained almost constant in mice (Fig. 4G, H). These observations indicate that the difference in tissue growth arises from cell volume regulation rather than from the regulation of cell number. Based on cell volume and cell number measurements, we estimated the cell volume ratio under the assumption that it is constant over time: *α* ≈ 0.5 in zebrafish and *α* ≈ 1 in mice. The theoretical tissue growth curve (tissue volume plotted as a function of cell number) corresponding to these values shows good agreement with experimentally observed growth curves, supporting consistency between the model and the data (Fig. 4E).

To verify whether the cell volume regulation constrains tissue growth across species, we estimated *α* during OTV expansion in 12 species. Although *α* values varied among species, the Z-mode species consistently exhibited *α* values close to 0.5, whereas the M-mode species showed *α* values of approximately 1 or greater (Fig. 4I, J). Across all examined species, *α* broadly corresponded to *β* (Fig. 4C, J), indicating that distinct modes of cell volume regulation underlie the diverse OTV expansion modes.

These results demonstrate that divergent OTV expansion modes across vertebrates operate within a shared mechanical framework. At the cellular level, stress sensitivity can be manifested through changes in cell number during OTV expansion (Fig. 4F, fig. S5C, F). Importantly, irrespective of stress-dependent changes in cell number, the mode of cell volume regulation determines how this cellular stress sensitivity is translated into tissue-scale volume responsiveness, thereby quantitatively linking diversity in OTV morphogenesis to cellular volume control.

### Heterochrony links cell volume regulation during organogenesis to embryo size and developmental tempo

Having established the mechanical basis of divergent OTV expansion processes, we next asked how these organ- and development-scale phenomena can relate to embryo- and evolution-scale developmental strategies. We first analyzed temporal changes in cell volume by tracking the average cell volume along the cell lineage leading to the inner ear, from cleavage to organogenesis, in zebrafish and mice. The comparison revealed similar decay profiles but distinct timings relative to the developmental timeline (Fig. 5A). In zebrafish, cell volume continued to decrease until the late OTV expansion phase, as if the cleavage-like state extended to the middle of the organogenetic period. In contrast, in mice, the cell volume reduction ceased by the end of cleavage—well before organogenesis—after which the cell volume remained constant (Fig. 5A). This difference in the developmental timing represents a consequence of heterochrony between zebrafish and mice.

**Fig. 5.**
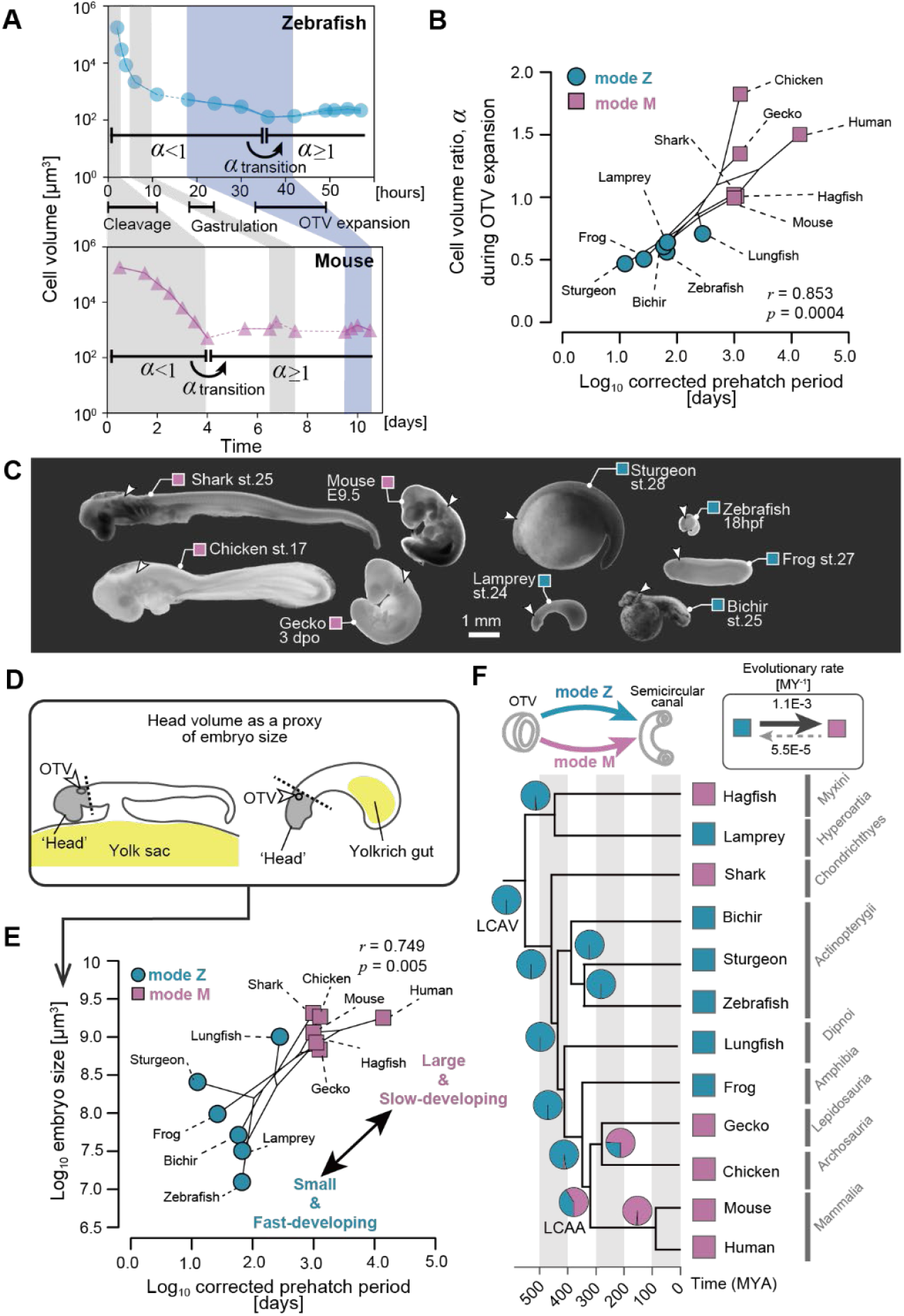
Subphylum-scale phylogenetic comparison and evolution of the morphogenetic process. (**A**) Mean cell volume at each developmental stage in zebrafish (top) and mice (bottom). Cell volume was calculated along the cell lineage leading to the inner ear. Data obtained from different studies are connected by dashed lines. Zebrafish data at 18–48 hpf and mouse data from E9.5–10.5 are from Fig. 4G. Other data are from refs. (*53-57*). Mean ± SE. (**B**) Correlation between *α* and the corrected developmental period. Phylogenetic canonical correlation analysis. (**C**) Left lateral view of whole mount images of fixed embryos from representative animals at around mid-developmental stages at the same magnification. Colored boxes beside animal names represent the mode of the process they adopt (Z-mode, blue; M-mode, magenta). Arrowheads show the position of the OTVs. (**D**) Schematic drawing representing the ‘head’ region (shaded in gray), arbitrarily defined in this study. (**E**) Log_10_ embryo size as a function of log_10_ prehatching developmental periods corrected by rearing temperature. Phylogenetic canonical correlation analysis. (**F**) Diversity of the OTV growth processes between mode Z (blue box) and mode M (magenta box) mapped on the tips of the vertebrate phylogenetic tree and ancestral states reconstructed by the M*k* transition model. Pie charts show the maximum likelihood at the nodes. The upper-right inset shows calculated evolutionary transition rates between traits. LCAA, last common ancestor of amniotes; LCAV, last common ancestor of vertebrates; MYA, million years ago. Scale bar = 1 mm (C).

Assuming that similar cell-size dynamics observed in zebrafish and mice are shared across the other examined vertebrate species, the relative timing of the onset of inner ear development with respect to the cell-size dynamics can be inferred from the value of *α* during OTV expansion (Figs. 4I and 5A). When *α* ≈ 0.5, OTV expansion begins while cell volume is still decreasing; when α ≈ 1, expansion starts after this reduction has ceased. Interspecies differences in α during OTV expansion thus reflect whether the α transition (from α ≈ 0.5 to ≥ 1) occurs before or after the onset of OTV expansion. Indeed, α values during OTV expansion in the Z- or M-mode species (Fig. 4I, J) indicate that Z-mode species initiate OTV expansion before completing the *α* transition, whereas M-mode species begin it afterward.

Given that the relative timing of the *α*-transition within the developmental program differs between Z-mode and M-mode species, we next examined how such differences in relative developmental timing relate to the overall developmental duration. To address this question, we analyzed correlations among the value of *α* during OTV expansion, developmental duration, and the mode of inner ear morphogenesis (Fig. 5B, fig. S7). Because ectothermic animals modulate their developmental rate depending on environmental temperature, we corrected the developmental duration for rearing temperature to obtain a temperature-standardized developmental duration for each species (Materials and Methods). Phylogenetic comparative analysis revealed that species with a smaller *α* during OTV expansion tend to have shorter developmental durations, whereas species with a larger *α* tend to have longer developmental durations (Fig. 5B). These results suggest that the modes of the inner ear morphogenesis reflect global embryonic features associated with the developmental strategy, rather than being restricted to a local, organ-specific phenomenon.

Previous studies have demonstrated a typical size-speed trade-off in embryos across animal taxa (*34, 35*) and that embryo size, alongside the developmental duration, is a salient indicator of embryonic trait diversity (*36*). We therefore tested whether embryo size correlated with the two modes of inner ear morphogenesis. We approximated embryo size by measuring the volume of the ‘head’ region (including the otic and preotic region) at the earliest stage of OTV expansion (Fig. 5C, D), allowing us to quantitatively compare embryo size while minimizing confounding effects arising from interspecific variation in yolk volume and somite number (*37, 38*). Phylogenetic comparative analyses revealed that Z-mode species in the inner ear morphogenesis tend to have smaller embryo size and shorter developmental periods, whereas M-mode species are characterized by larger embryo size and longer developmental duration (Fig. 5E; fig. S7). These results indicate that the embryo type—small and fast-developing versus large and slow-developing—is correlated with the mode of inner ear morphogenesis (Fig. 5C, E).

Finally, to place the divergent modes of inner ear morphogenesis in a context of evolutionary history, we reconstructed their ancestral states in the vertebrate phylogeny. We assumed that the Z- or M-mode of inner ear morphogenesis is a binary trait, that either mode is plesiomorphic in vertebrates, and that evolutionary transitions between the two modes are possible. The best-supported ancestral reconstruction estimated that the last common vertebrate ancestor adopted the Z-mode inner ear morphogenesis, followed by at least three independent transitions to the M-mode in descendant lineages (Fig. 5F). The estimated evolutionary transition rate from mode Z to M was approximately 20-fold higher than the reverse direction, indicating a pronounced asymmetry in directional evolution (Fig. 5F, inset).

## Discussion

Our integrative evolutionary developmental biology (Evo-Devo) and tissue mechanics approach revealed cryptic diversity in the mechanics of vertebrate inner ear morphogenesis. We identified two distinct modes of OTV expansion: mode Z, characterized by constant epithelial volume, and mode M, characterized by constant epithelial tissue stress. These mechanical modes correlate with the type of embryos defined based on embryo size and developmental tempo (Fig, 5E, fig. S8). We propose that the evolutionary transition from Z- to M-mode inner ear morphogenesis reflects a heterochronic shift; delaying the onset of organogenesis permits epithelial volumetric growth after *α* transition and is accompanied by an adaptive shift of embryonic traits from small, fast-developing embryos to large, slow-developing ones (fig. S8).

The evolutionary shifts in inner ear morphogenesis from mode Z to M (Fig. 5F) likely are promoted under reproductive environments that favor large embryos with a prolonged developmental period, such as deep-sea environments, oviposition on land, or exposure to high predation pressure (*1, 39*). Embryos of M-mode species use stress-dependent epithelial growth presumably to ensure tissue integrity despite their larger size and prolonged developmental period. M-mode morphogenesis can be time-consuming because the pace of morphogenesis is constrained by the length of each cell cycle. In contrast, embryos of Z-mode species complete OTV expansion largely within the window of early tissue-growth arrest, presumably to ensure rapid morphogenesis. These alternative mechanics consequently buffer the seemingly conflicting demands imposed by divergent developmental strategies, enabling embryos of different sizes and developmental tempos to achieve primordium expansion through mechanically distinct means, while ultimately ensuring the structural and functional reproducibility of the inner ear.

The apparent conflict between the diversification of embryonic traits associated with distinct developmental strategies and the preservation of ancestral organ structure as seen in the inner ear can be viewed as a manifestation of opposing selective pressures. Embryo size or developmental duration is likely shaped by directional or frequency-dependent selection (*40*), whereas the morphogenetic process of the inner ear is presumably under stabilizing selection to maintain its structural and functional outcome. As discussed above, heterochronic shifts in the relative timing of the *α* transition mechanistically link embryo size, developmental duration, and the divergent inner ear morphogenetic processes in development (fig. S8), making this timing sensitive to both selective regimes in evolution. Thus, just as theoretical studies predict that the evolution of DSD is driven by genetic factors that simultaneously regulate multiple traits exposed to opposing selective pressures (*12, 13*), the timing of the *α* transition may likewise act as a ‘pleiotropic factor’ underlying the evolution of DSD in the inner ear morphogenesis.

Our findings highlight two intriguing factors that shape species-specific inner ear morphogenetic processes. The first is mechanosensitive tissue growth, in which proliferation depends on tensile stress, likely involving mechanosensitive molecular pathways described previously (*41*). The second is tissue-growth arrest driven by reductions in cell volume, likely reflecting the relative timing of the proposed *α* transition, for which the timing in turn would be mechanistically under the regulation of some molecular and mechanical cues (*42, 43*). The volume-reducing divisions observed during organogenesis in fast developers resemble those seen during the cleavage stage in early development. However, whether the molecular or cellular mechanisms reported from studies examining cleavage (*44*) also operate at these much later stages of development remains unaddressed in the present study. Comparable growth arrest during organogenesis may not be unique to inner ear morphogenesis and may appear in other organ systems. For instance, during eye morphogenesis, mice and humans require cell proliferation for optic cup formation (*33, 45*), whereas zebrafish can form optic cups even when cell division is blocked (*46, 47*). Likewise, during axial elongation, transient volumetric growth in the tailbud mesenchyme occurs in sharks and mice but not in lampreys or zebrafish (*48*). These examples suggest that the distinction of Z-mode (less dependent on tissue growth) versus M-mode (driven by tissue growth) morphogenesis may apply to organ development beyond the inner ear.

Taken together, our study advances the emerging framework of Evo-Devo mechanics by elucidating a key mechanical principle that underlies divergent processes yielding conserved morphology. Progress in this field relies on quantitative, cross-species analyses that can distill complex morphogenetic behaviors into simple and common geometric and mechanical parameters. In our study, we integrated exhaustive measurements with simple mathematical formulations to develop a mechanical framework of inner ear morphogenesis in zebrafish and mice and to extrapolate this framework to a broader range of vertebrate taxa. This approach allowed us to capture essential species-specific differences in inner ear mechanics, such as the cell volume ratio *α*, and to discuss potential selective pressures that have driven the direction of evolution. Such quantitative approaches—combining quantitative description, experimental perturbations, mechanical inference, and minimal modeling—provide a common analytical language for comparing divergent developmental processes and for revealing the mechanical principles that underlie their evolution (*24, 49-51*). Future research may benefit from applying and refining such quantitative approaches to elucidate the mechanical basis of Evo-Devo (*52*).

## Acknowledgements

We thank the members of the Kamei lab for their maintenance of the zebrafish aquarium at Kanazawa University; S. Terashima and A. Matsuoka for their assistance with 3D data processing; the staff of Fujikin Corporation for providing sturgeon embryos; M. Yamashita for her assistance with collecting bichir embryos; the members at RIKEN LARGE for providing gecko embryos; D. T. Roberts for his assistance with collecting lungfish embryos; M. Hoshino and K. Uesugi for their assistance with SRXµCT; K. G. Ota and Y. Oisi for their pioneering works using hagfish embryos at Kuratani lab; the Virtual Human Embryo Project, the 3D Atlas of Human Embryology, and the Human Developmental Biology Resource for sharing high quality histological data of human embryos with the science community; and Y-C. Wang and Y. Morishita and their lab members for valuable feedbacks and discussions. The *Xenopus laevis* albino strain was provided by Hiroshima University Amphibian Research Center (RRID: SCR_019015) through the National BioResource Project (NBRP) of MEXT, Japan. The authors used ChatGPT 5.2 to improve the language and readability of the manuscript.

## Funding

This work was supported by the following:

The Japan Society for the Promotion of Science (JSPS) KAKENHI (Grant Nos. 23K19363, 25K18522, and 25KJ0149 to S. Kuroda),

JSPS KAKENHI (Grant No. 24KJ0090 to S. A. H.),

JSPS KAKENHI (Grant No. 23K19369 to S. Kato),

JSPS KAKENHI (Grant Nos. 22H0517, 24H01398, 24H01937, 25K01118, and 25K22468 to S. O.),

The Japan Science and Technology Agency (JST) CREST (Grant Nos. JPMJCR1921 and JPMJCR24B2 to S. O.),

JST PRESTO (Grant No. JPMJPR25KB to S. A. H.).,

The Japan Agency for Medical Research and Development (AMED) Program for Technological Innovation of Regenerative Medicine (Grant No. 23bm0704065h0003 to S. O.),

The World Premier International Research Center Initiative (WPI), Ministry of Education, Culture, Sports, Science and Technology (MEXT), Japan (to S. O.).

## Author contributions

Conceptualization: S.Kuroda, S.A.H., S.O.;

Funding acquisition: S.Kuroda, S.A.H., S.Kato, S.O.;

Experiments: S.Kuroda, S. Kato, W.T., T.H., K.F., K.H., N.S., A.Y.;

Supervision: S.O.;

Visualization: S.Kuroda., S.A.H.;

Writing – original draft: S.Kuroda, S.A.H., S.O.;

Writing – review & editing: S.Kuroda, S.A.H., S.Kato., O.H., H.K., W.T., S.H., T.H., K.F., M.S., H.O.,

H.K., S.Kuratani, and S.O.

All authors confirmed the final version of the manuscript.

## Data and materials availability

All raw imaging data and analysis code will be available at the SSBD:database upon publication.

## List of Supplementary Materials

### Supplementary Materials

Materials and Methods

Figs. S1 to S8

Tables S1 to S4

Movie S1 (separate file)

Data S1 to S4 (separate files)

## Supplementary Materials

### Materials and Methods

#### Embryos

All experiments were performed in compliance with the experimental designs approved by Institutional Animal Care and Use Committee at Kanazawa University (protocol#: 6-2757, 6-2398, AP24-017, AP24-042, AP24-067, AP24-72), and RIKEN (protocol#: A2001-03-94). Adult lampreys (*Lethenteron camtschaticum*) were kept in the aquarium at the University of Tokyo at 12°C until sexual maturation. Lamprey embryos were obtained via artificial fertilization and raised in the 10% Steinberg solution [5.82 mM NaCl, 0.067 KCl, Ca(NO_3_)_2_ 4H_2_O, 0.083 mM gSO_4_ 7H_2_O, 0.5 mM HEPES] containing 0.6 ppm Methylene Blue (Cat.#:19-3240, Sigma-Aldrich) at 9–18 °C. Adult sharks (*Scyliorhinus torazame*) were kept in the water tank at the University of Tokyo at 16°C. Shark embryos were obtained via natural fertilization and raised in the sea water containing 0.2% antibiotic-antimycotic mixed solution (Cat.#: 09366-44, Nacalai Tesque) at 16 °C. Adult bichirs (*Polypterus senegalus*) were kept in the aquarium at Niigata University at 28 °C. Bichir embryos were obtained via natural fertilization and raised in the embryo medium (13.7 mM NaCl, 0.54mM KCl, 0.025 mM Na_2_HPO_4_, 0.044mM KH_2_PO_4_, 1.3mM CaCl_2_, 1.0 mM MgSO_4_, and 0.42 mM NaHCO_3_) at 23-28.5°C. Adult sturgeons (hybrid of *Huso huso* and *Acipenser ruthenus*) were bred in the hatchery of the Fujikin Corporation. Fertilized eggs were collected in the breeding season on March to April and raised in the embryo medium at 15 °C. Adult wild-type zebrafish (*Danio rerio*, AB strain) were provided by the Zebrafish International Resource Center (Oregon, USA) and were kept at 28.5°C on a 14-hour night/10-hour dark cycle in the aquarium at Kanazawa University. Zebrafish embryos were obtained via natural fertilization and were kept in the embryo medium at 28.5°C. Fertilized lungfish (*Neoceratodus forsteri*) embryos were collected from a breeding facility in Cooroy (Queensland, Australia) under the permission of the General Fisheries Permit (GFP 209221) from the Queensland Government. Fixed samples were transported to Japan under the permission of Convention of International Trade in Endangered Species of Wild Fauna and Flora (CITES, certificate#:PWS2023-AU-003043). Adult albino (*Xla*.*tyr*^*emOgino*^) frogs (*Xenopus laevis*) were originally generated with CRISPR/Cas9 targeted mutations in tyrosinase genes in the previous study (*58*) and bred in Hiroshima University. Embryos were raised in the 0.3x MMR (10x MMR: 1 M NaCl, 20 mM KCl, 20 mM CaCl_2_ 2H_2_O, 10 mM MgSO_4_, 50 mM HEPES) by supplementing 10 μg/ml gentamicin (Cat.#: 078-06061, Fujifilm) at 14-22 °C. Adult gecko (*Paroedura picta*) colony were kept in the Laboratory for Animal Resources and genetic Engineering (LARGE), RIKEN at 26–28 °C. Eggs obtained by natural fertilization were incubated in the humidified laboratory chamber at 28 °C. Chicken (*Gallus gallus*) eggs were purchased from a local farm and incubated in the humidified chamber at 38 °C. Adult albino (Slc:ICR) mice (*Mus musculus*) were bred in the colony at Sankyo Labo Service Corporation and pregnant female mice were shipped to the laboratory at Kanazawa University. Mice embryos were staged according to passage date, with midnight on the day the plug was identified as E0. Fertilized eggs of lampreys, sharks, bichirs, sturgeons, frogs, and geckos were shipped to the laboratory at Kanazawa University and raised in the lab incubator to the desired stages. Except for mice, embryonic stages were determined based on external morphology, with reference to staging tables; lamprey–ref (*59*), shark–ref (*60*), bichir–ref (*61*), sturgeon–ref (*62*), zebrafish–ref (*21*), lungfish–ref (*63*), frog–ref (*64*), gecko–ref (*65*), and chicken–ref (*66*). Histological data from hagfish (*Eptasterus burgeri*) and human (*Homo sapiens*) embryos were obtained from specimens prepared in the previous research (*17*) and open data resources, including the 3D Atlas of Human Embryology (3AHE; https://www.3dembryoatlas.com/), Virtual Human Embryo (VHE; https://virtualhumanembryo.lsuhsc.edu/), and Human Developmental Biology Resource (HDBR; https://hdbratlas.org/). These databases of human embryology are available for use in non-profit research purposes.

#### Fixation and imaging

Embryos were fixed by Bouin’s fixative (mixture of saturated picric acid, 37% formaldehyde, and acetic acid at a volume ratio of 15:5:1) or Serra’s fixative (mixture of 100% ethanol, 37% formaldehyde, and acetic acid at a volume ratio of 6:3:1) overnight at 4 °C and stored in 70% ethanol until use. Embryos of aquatic species were anesthetized by adding 0.17 mg/ml ethyl m-aminobenzoate methanesulfonate (MS222, Cat.#: E10521, Sigma-Aldrich) in the incubation media just before fixation. Before histological preparations, bichir embryos were bleached by incubating in 100% methanol containing 5% hydrogen peroxide (Cat.#: 081-04215, Wako) under light exposure for one to three days while monitoring the pigment residues.

For whole-mount tissue staining, embryos were rinsed in phosphate-buffered saline containing 1% triton-X (PBS1TX) three-times and transferred in 2.5 μg/ml rhodamine B isothiocyanate (RBITC; Cat.#: 283924, Sigma-Aldrich) in PBS1TX for one-to-seven overnights at room temperature depending on specimens’ sizes. Stained samples were washed by PBS1TX for more than five times to remove excess dye, followed by tissue clearing using CUBIC-R reagent (Cat.#: CSCR-002, CUBICStars) according to the manufacturer’s protocol. Instead of CUBIC method, some specimens rich in yolk platelets in somatic cells (e.g., lamprey, bichir, sturgeon, frog embryos) were dehydrated with 100% methanol and then cleared by transferring in the BABB solution (a 1:2 mixture of benzyl alcohol and benzyl benzoate). Cleared samples were set in a glass bottom dish and imaged by LSM800 (Zeiss) or FV3000 (Olympus) inverted confocal microscopes equipped with objective lenses [Plan-Apochromat 10x/0.45 M27, LD LCI Plan-Apochromat 25x/0.8 Imm Korr DIC M27 (Zeiss), UPLXAPO10X, UPLXAPO20X, or UPLXAPO30XS (Olympus)].

For preparing histological sections, embryos were dehydrated through a graded ethanol series, methyl benzoate, and benzene, then embedded in paraffin (Cat.#: P3683, Sigma-Aldrich) melted at 65°C. Sections were cut at a thickness of 6–10 μm using a microtome (Cat.#: HM 335E, Microm). After deparaffinization, sections on a slide were stained with hematoxylin and eosin following standard protocols.

For lungfish imaging, paraffin-embedded embryos were scanned using high-resolution X-ray tomography (voxel size = 2.26 μm, energy = 25 keV) in the experimental hatch of the beamline BL20B2 at the SPring-8 synchrotron radiation facility in Sayo-Cho, Hyogo Prefecture, Japan. X-ray images were converted to visible light using a 500 µm-thick Lu_3_Al_5_O_12_:Ce+ (LuAG) scintillator, with a sample-to-detector distance of 1000 mm. A total of 1800 projections over 180° were acquired at 300 ms exposure per projection. Tomographic reconstruction was performed using a filtered back-projection algorithm (*67*).

The fixation, staining, clearing (if applicable), and imaging methods for each sample are detailed in data S1.

#### Measurement of luminal pressure

The lab-made pressure measurement device was designed and assembled inspired by previous works (*27, 68*). Briefly, a piezo-resistive silicon pressure sensor (Cat.#: HSCDANT001PGSA3, Honeywell) was connected with a glass capillary via a polytetrafluoroethylene tube (inner diameter (ID), 1 mm; outer diameter (OD), 2 mm). A link between the capillary and connecting tube was sealed with a silicon tube (ID, 0.8 mm; OD, 4 mm). When assembling the system, a sensor, capillary, and connecting tubes were filled with deionized water while keeping air bubbles from getting inside. Glass capillaries were prepared using a vertical puller machine (PC-10, Narishige) with the following setting: heat1 = 55, heat2 = 50, adding further weight of 360 g). The tip of the needle was broken using forceps to obtain ∼35 um tip OD. The inner surface of the glass needle was coated with NovyCoat (Cat.#: FG-NVC100, FastGene) prior to the measurement to prevent clogging by tissue debris. For the data collection, the pressure sensor was connected to a microcontroller board (Cat.#: UNO R3, Arduino). Measurements were collected via a sensor evaluation board (Cat.#: SEK001, Honeywell). Data were acquired at a sampling rate of 5 Hz and exported as .csv format. Circuit connections were followed according to the manufacturer’s instruction (https://sps.honeywell.com/us/en/products/advanced-sensing-technologies/healthcare-sensing/sensor-evaluation-kits/).

For sensor calibration, we submerged the glass needle and recorded the hydrostatic pressure at different depths in the water column. After plotting measured pressure as a function of submerged depth, we performed the linear regression by using statsmodels (v. 0. 14. 2) (*69*) library in Python (v. 3. 10. 11) (fig. S4B). The estimated slope of the regression line was 9.69 ± 0.04 (SE) Pa/mm (*R*^*2*^ = 0.967), which is comparable to the expected value of 9.81 Pa/1 mm H_2_O.

Mouse embryos were secured between trenches made of 2% agarose gel in PBS during measurement. The pressure of the OTV explants was measured in the collagen gel. A glass needle connected to the pressure sensor was inserted into the OTV, and the needle position was held steady for six minutes and retracted (figs. S4C, S5G, H). The baseline pressure was calculated as the average value during one minute prior to needle insertion. The inner ear pressure was measured as the difference between this baseline and the average value during the one minute prior to needle retraction. The calculation for determining the measured values was performed using NumPy (v. 1. 24. 3) (*70*) library in Python. All measurements were listed in data S2.

#### Mouse explant culture

Mouse embryos at E9.5 were isolated from a pregnant mouse and collected in a dish filled with D-MEM containing 0.5X penicillin-streptomycin at 37°C. Isolated embryos were then placed on a gel plate (2% agarose in PBS) filled with PBS and OTVs were dissected along with their surrounding tissues using sharpened tungsten needles under a stereomicroscope (SZX16, Olympus). OTV explants were collected into the 1.5 ml tube filled with the explant culture medium [10% Knockout serum replacement (Cat.#: 10828010, Gibco), 1% N-2 supplement (Cat.#: 17502048, Gibco), 2% fetal bovine serum (Cat.#:12483020, Gibco), and 0.5x penicillin-streptomycin in D-MEM] and embedded in 240 µl of collagen gel on the glass bottom dish. After waiting for solidifying of the gel, the dish was filled with 3 ml of culture medium and cultured for one day under 37 °C, 5 % CO_2_, 90%< (relative humidity) environment.

#### Pharmacological treatments

For pharmacological treatments in zebrafish embryos, dechorionated zebrafish embryos at 18 hpf were incubated in the embryo medium containing 1% dimethyl sulfoxide (DMSO; for mock treatments) or 1 mM ouabain (Cat.#: 1076, Tocris) (inhibition of fluid transport) or 150 µM aphidicolin (Cat.#: A0781, Sigma-Aldrich) (inhibition of cell division) for 12 hours. For pharmacological treatments in the mouse OTVs, mouse OTV explants were incubated in the explant culture medium containing 1 % DMSO (mock) or 0.1 mM ouabain or 0.5 µM aphidicolin for 24 hours. Embryos and explants were anesthetized in MS-222 (only applicable for zebrafish) and fixed by Bouin’s fixative for volume measurements (data S3).

#### Volume segmentation and morphometric analysis

For 3D segmentation and quantification, generated serial confocal images were converted into serial tiff image using Fiji/ImageJ (v. 1.54g, NIH), imported into Drishti software (v. 3.2, https://github.com/nci/drishti), and exported as pvl.nc format. In Drishti, tissue and lumen area were separately segmented out based on tissue histology visualized by RBITC staining or sample green autofluorescence coming from Bouin’s fixative. Tissue and lumen volumes were calculated using the software function. Segmented volumes were subsequently converted into 3D mesh files in ply format and surface areas were calculated in ParaView (v. 5.12.0). 3D reconstructed images were visualized by rendering ply files in Blender (v. 4.0.2). Measurements were collected in Microsoft Excel and plotted using Matplotlib library (v. 3.7.1) in Python.

#### Protrusion shape in the semicircular canal formation

The timing when the protrusions extending from both poles of the OTV contacted each other as a fusion plate was captured in 2D (confocal or histological) sections, either from previous studies or newly acquired images. Using the ‘Straight’ ROI tool in Image J/Fiji, the depth (*d*_*1*_, *d*_*2*_) and base width (*w*_*1*_, *w*_*2*_) of protrusions were measured (fig. S2A). Based on these measurements, the shape of the protrusion ω, defined as 2(*d*_1_ + *d*_2_)/(*w*_1_ + *w*_2_), was calculated. Measurements are deposited in table S1.

#### Normalization of time and geometry

For all animal species investigated in this study, OTV volume increased monotonically during the OTV expansion process (fig. S1). We therefore introduced pseudotime τ at a given developmental stage *t* by the non-dimensionalized changes in OTV radius to align corresponding stages and enable cross-species comparisons, namely,

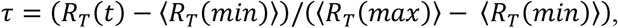

where ⟨*R*_*T*_(*mix*)⟩ and ⟨*R*_*T*_(*max*)⟩ are average total OTV radii at the earliest and latest stages of OTV expansion (fig. S1), respectively.

To compare the deformation of the OTV during the expansion at τ (0 ≤ *t* ≤ 1), strains in epithelial height was expressed as

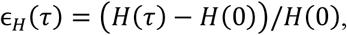

where *H*(τ) is epithelial height at τ.

For a comparison of the balance between lumen and tissue volume changes during OTV expansion across species, we normalized lumen and tissue volumes changes as

*Norm.log lumen volume change* = *log* (*V*_*L*_)/*V*_*L*_(0))/*log*(*V*_*T*_(1)/*V*_*T*_(0)), and *Norm. log tissue volume change* = *log*(*V*_*E*_(*t*)/*V*_*E*_(0))/*log*(*V*_*T*_(1)/*V*_*E*_(0)), respectively, where *V*_*L*_(*t*) and *V*_*E*_(*t*) are lumen and tissue volume at a given developmental stage *t, V*_*L*_(0) and *V*_*E*_(0) are average lumen and tissue volumes of at τ = 0, and *V*_*T*_(0) and *V*_*T*_(1) are average total OTV volume at τ = 0 and 1, respectively.

#### Estimation of cell number and volume

To estimate the total number of constituent cells in the OTV, we counted the number of cells within a local area, using confocal/histological sections at the level near the basal surface of the lateral pole of OTV, and then calculated the average number of cells per unit surface area (mm^-2^). By multiplying this value by the total surface area of the OTV, we estimated the total number of constituent cells. The average cell volume was determined by dividing the epithelial volume by the estimated number of cells.

Measurements of cell volumes from non-OTV lineages were adapted from the following references: zebrafish—ref.(*53, 57*) and mice—refs.(*54-56*). The measurements used for the calculations are provided in data S1 and S4.

#### Estimation of the mechanical state of the OTV

Assuming the geometry of OTV as a spherical shell with inner (lumen) radius *R*_*L*_, and outer (total) radius *R*_*T*_, the lumen, epithelial tissue, and total volumes are given by 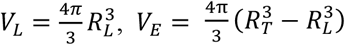 and 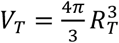 respectively. We assume that the OTV is in a mechanical equilibrium at each developmental timepoint and the thickness (cell height) *H* = *R*_*T*_ − *R*_*L*_ is relatively small compared to the radii. Young–Laplace equation (*30*) describe the force balance relationship between the hydrostatic pressure *P* in the lumen and the effective tensile stress *σ* in the circumferential direction as

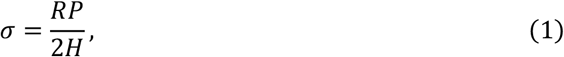

where *R* is a representative radius and we set *R* = *R*_*L*_ for simplicity. Note that *P* should be interpreted as the difference between the pressure acting on the inner surface and that on the outer surface. The pressure on the outer surface is assumed to be zero in the following calculations. Although the surrounding tissue may exert pressure on the outer surface of OTV, it is difficult to estimate the value, and a constant external pressure does not qualitatively change the conclusion.

The circumferential stress σ was estimated using the above equation. Since the measurement timings of the volumes and the pressure are different, linear interpolation and extrapolation were performed on the pressure data to estimate the pressure values at the times of geometry measurements. The uncertainty of the estimate was calculated by the bootstrapping method, and 95% confidence intervals are shown in the figures (Figs. 2F, 3K). The statistical hypothesis test in Fig. 3K was based on the bootstrap resampling (N=10000) (*71*), where the null hypothesis was that the two groups have the same median circumferential stress.

#### Dynamical modeling of the OTV expansion

Our mathematical model of OTV is composed of a lumen filled with incompressible fluid and epithelial tissue with elastoplastic properties. The fluid flux into the lumen is assumed to be independent of tissue growth and mechanical changes and proportional to the lumen surface area

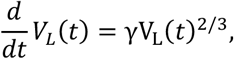

where *γ* > 0 corresponds to the fluid flux per unit area. The lumen inflation stretches the epithelial tissue in the circumferential direction and the epithelial tissue generates a tension in response to the stretching, whose tensile stress σ is assumed to be given by

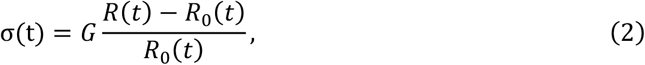

where *G* is the effective stiffness, *R* is a representative radius, which is interpreted to be lumen radius *R* = *R*_*L*_, and *R*_0_ is the corresponding reference length. At the timescales of our observation, we assume the tissue growth modifies the reference radius *R*_0_, which could relax the stress in Eq. (2). To explain the observed constancy of the stress in mouse OTVs, we hypothesize that tissue growth rate is proportional to the deviation of the current stress σ from the constant target stress σ^∗^ so that the epithelial tissue volume satisfies

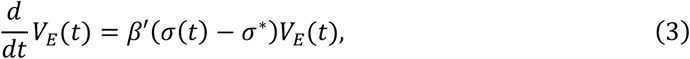

and that the reference radius is given by

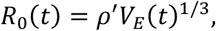

where *β* = *β*^′^*G* is the tissue growth rate coefficient and 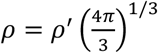 is the shape parameter that corresponds to the tissue-to-lumen volume ratio in the stress-free configuration. One can obtain the analytical solution of the dynamical system as

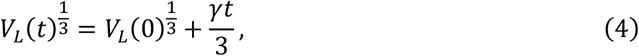

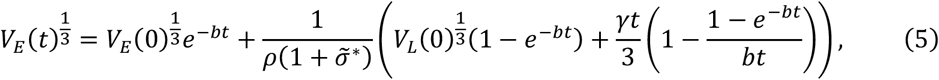

where 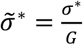 and 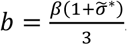. If *β* = 0, the tissue does not grow *V*_*E*_ (*t*) = *V*_*E*_ (0) and the stress increases linearly in time, 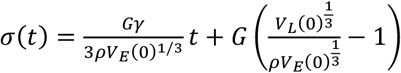, as observed in zebrafish. If *β* > 0, the tissue volume and the stress behave like 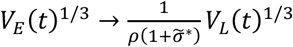 and σ(*t*) → σ^∗^ at long timescales (*t* → ∞) reproducing the isometric scaling and stress constancy in mouse.

To quantitatively verify the agreement of the mathematical model with the observed developmental processes of mice and zebrafish, we performed the Markov chain Monte Carlo sampling of the model parameters. The length scales in the model were nondimensionalized by setting empirical average of the initial tissue volume to be 1. Among six parameters (*γ, β, V*_*L*0_, *ρ, G*, σ^∗^) to be estimated, the three (*γ, β, V*_*L*0_) were treated as species-specific parameters and the other three (*ρ, G*, σ^∗^) were shared parameters. Due to the limitation of the data, the introduction of shared parameters was necessary to identify parameters. The prior distributions were *log G ∼ N* (0,20^2^), *γ ∼ Exp* (10), 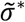, *ρ β ∼ U* (0,1), and 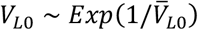, where 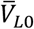 denotes the empirical average of *V*_*L*0_. The observation distributions for *V*_*L*_, *V*_*E*_, and *P* were Gaussian distributions with mean given by Eqs. (4) and (5). The standard deviations were 0.1 for *V*_*L*_ and *V*_*E*_, and 10 [Pa] for zebrafish and 50 [Pa] for mouse. Since we had no pressure data for early phase of zebrafish OTV expansion, we assumed *P*(0) ∼ *N*(0, 0.1^2^). The almost zero pressure, equivalently almost zero stress, is reasonable because the lumen volume is close to zero at this stage. We used No-U-Turn Sampler (*72*) implemented in numpyro (*73, 74*) with 2500 samples with 500 warmup.

We investigated whether the same mathematical model could be applied to 12 vertebrate species, including mice and zebrafish. However, information on pressure *P* and physical time *t* was not available, and only information on the volumes of the lumen and tissue at the sampled developmental stages was available. Therefore, we performed fitting in a different way from that used for mice and zebrafish. First, the three parameters (*ρ, G*, σ^∗^) were fixed to the posterior expectations obtained from the fitting for zebrafish and mice. Although there is no evidence to suggest that these parameters are common across species, this assumption allowed us to identify the parameters, and we confirmed that the mathematical model fit the data well under this assumption. Second, we inferred the times *t* of observations from the lumen volumes *V*_*L*_. As can be seen in Eq. (4), the change in the cube root of the lumen volume is directly proportional to the elapsed time *t*. Thus, we arbitrarily set *γ* = 1 and estimated the time *t*_*i*_ of *i*-th measurement as *t*_*i*_ = 3V_*L*_(*t*_*i*_)^1/3^ − *V*_*L*_(0)^1/3^. The prior distributions were Δ*t*_*i*_ ∼ *Ex*p(1) and β ∼ *U*(0, 5), where Δ*t*_*i*_ := *t*_*i*_ − *t*_*i*−1_ denotes the time interval between consecutive observations. Other prior and observation distributions were the same as in the sampling for zebrafish and mice.

Note that the estimated β for human was as small as that for other Z-mode species, even though human is clearly classified as mode M in terms of thickness change, protrusion shape, and cell volume ratio α (Figs. 1C, 1E, 4I). The deviation is presumably due to the high expansion rate of both lumen and tissue in the human OTV. The assumption of shared mechanical parameters (*ρ, G*, σ^∗^) was necessary for parameter estimation, but it does not generally hold. Although direct mechanical measurements of the human OTV expansion are inaccessible, species-specific measurements of these mechanical parameters would resolve the discrepancy.

#### Calculation of cell volume ratio

Denoting the epithelial tissue volume as *V*_*E*_(*t*) and the number of cells in the tissue as *N*(*t*), the average volume of a cell at time t is given by *v*(*t*) = *V*_*E*_(*t*)/*N*(*t*). Let us derive the dynamics of the average cell volume in the simplest case where the rates of population growth and volumetric growth are constant. Since the cell number doubles after average cell cycle length *φ*, the number of cells increases exponentially as

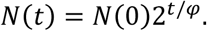

The cell volume is assumed to increase by a factor of alpha per cell cycle on average, thus we have

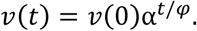

Taking the logarithm of base 2 on both sides of the above equations and eliminating *t*/*φ*, we obtain a formula to calculate α from the fold change of cell number and cell volume as follows:

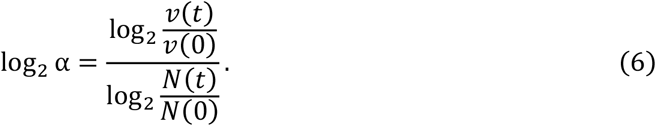

Alternatively, α can be interpreted as an index quantifying the coupling between population growth and tissue volumetric growth. Using the chain rule and 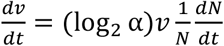, the time derivative of the tissue volume is given by

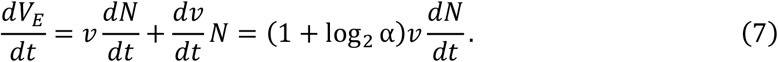

Thus, the tissue volume increases by (1 + log_2_ α)*v* for every new cell. When α = 1, we have 1 + log_2_ α = 1, meaning that volumetric growth is fully coupled to population growth. The tissue volume increases by the cell volume multiplied by the increase in cell number. When α = 0.5, we have 1 + log_2_ α = 1 − 1 = 0; there is no volumetric growth, and tissue growth is completely decoupled from population growth. When 0.5 < α < 1, the two growth processes are only partially coupled. The volumetric growth is possible but inefficient; the cell proliferation rate must be 1/(1 + log_2_ α) times higher than in the fully coupled case to achieve the same amount of tissue growth. Finally, by comparing Eqs. (3) and (7), we obtain the relationship between α and *β*:

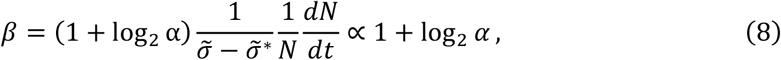

when 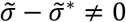

The value of α was estimated using Eq. (6) with medians of observed cell volumes and numbers. The uncertainty of the estimate was calculated by the bootstrapping method, and 95% confidence intervals are shown in the figures (Fig. 4).

#### Ancestral state reconstruction and phylogenetic comparative analysis

Ancestral state reconstruction for the discrete character was performed in R (v. 4.4.1). A time-calibrated vertebrate phylogenetic tree was obtained from TimeTree (https://timetree.org/) in Newick format. Because some of our target species were not directly available in the TimeTree dataset, we substituted them with closely related species: *Scyliorhinus torazame* was represented by *Scyliorhinus canicula*, and the hybrid *Huso huso* × *Acipenser ruthenus* was represented by *Acipenser ruthenus*. The trait matrix was constructed from mode of developmental processes (mode Z or M), coded as a binary state. We employed a Markov model and model fitting was performed using fitM*k* function in the phytools (v. 2.3.18) (*75*) package. The best-fitting model was selected based on the highest log-likelihood value. The posterior probabilities of ancestral states were visualized using pie charts at internal nodes.

To assess the evolutionary correlation between continuous characters while accounting for phylogenetic relationships, we applied phylogenetic canonical correlation analysis using phyl.cca function implemented in phytools package. The correlation strength *r* was estimated without fixing the lambda parameter, allowing the phylogenetic signal to be optimized. For visualization, plots were connected according to phylogenetic relationships using phylomorphospace function. To investigate the correlation between the mode of morphogenetic process (as a discrete trait) and a continuous trait while accounting for phylogenetic relationships, we performed phylogenetic correlation analysis using a Bayesian threshold model. The analysis was implemented by using the threshBayes function in the phytools. The threshBayes function defines prior distributions for model parameters as follows: the variances follow an exponential prior with a mean specified as 1000, the ancestral states follow normal priors with mean and variance specified as 0 and 1000, respectively, and the correlation coefficient *r* follows a uniform prior with a mean set to 0 over the interval [-1, 1]. We ran MCMC sampling with 100,000 generations and got the posterior distribution of the correlation efficient *r*.

#### Developmental period corrected by temperature

To calculate the prehatch embryonic period corrected by rearing temperature, we adopted a previously proposed model that describes relationships among developmental period *θ* in day, body mass at hatching *m*, and rearing temperature *T*_*c*_ in Celsius degree (*35*);

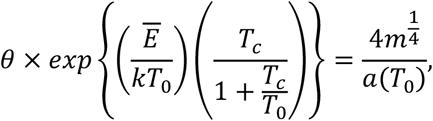

where *Ē* is the average energy for the reaction, *k* is Boltzmann’s constant, *T*_0_ = 273 K at which water freezes and biological reactions are assumed to cease, and *a*(*T*_0_) is a fundamental cellular biochemical property. Gillooly et al. showed that the observation data obtained from various animals spanned from ectotherms to endotherms, including fish, amphibians, and birds, can be fitted well using the equation (*35*). We assumed the relation is also applicable in animals examined in the present study. Thus, by substituting fitting parameters obtained in the Gillooly et al., namely 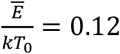 and 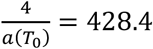, we obtain

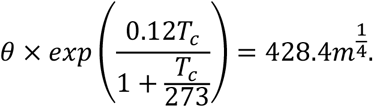

By putting 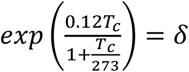, we utilized *δ* to correct developmental duration in the literature by rearing temperature *T*_*c*_ and get the corrected prehatch embryonic period as *θδ*. The data used to calculate *θδ* were listed in table S4 with references.

#### Geometry of semicircular canals

To evaluate the geometrical diversity of the functional inner ear in vertebrates, we parameterized the relative thickness of the anterior semicircular canal. We approximated the canal opening as an ellipse and obtained the major axis *D*_*M*_ and minor axis *D*_*m*_, as well as the canal thickness *d*, from the previously published data. Using the measurements, we calculated the ratio 2*d*/(*D*_*M*_ + *D*_*m*_) (fig. S7, table S3). The ellipse fitting was performed by manually segmenting the contour of the anterior semicircular canal opening using the Freehand Selection Tool in Fiji/ImageJ, followed by applying the Fit Ellipse function.

#### Statistics

No statistical methods were used to predetermine sample sizes but our sample size are similar to those reported in previous publications (*27*). In the pressure measurement, data were excluded if no pressure increase was recorded after needle insertion or if the measurements did not return to the baseline after needle removal. 3D segmentation was manually performed by a person distinct from an analyst to ensure blinding and avoid bias in data analyses. The boxplot displayed in the figures represents the interquartile range (IQR) with the median, while whiskers extended to 1.5 times the IQR. Data points that exceed 1.5 times the IQR above the third quartile or below the first quartile were considered outliers, yet outliers were included in the statistical analyses. Before comparing differences among experimental conditions, the normality test was performed using the Shapiro-Wilk test using the scipy.stats library (v. 1.10.1). Measurements were calculated with the function of Microsoft Excel (v. 2501) and with Pandas library and plotted with Matplotlib library (*76*) in Python.

**Fig. S1.**
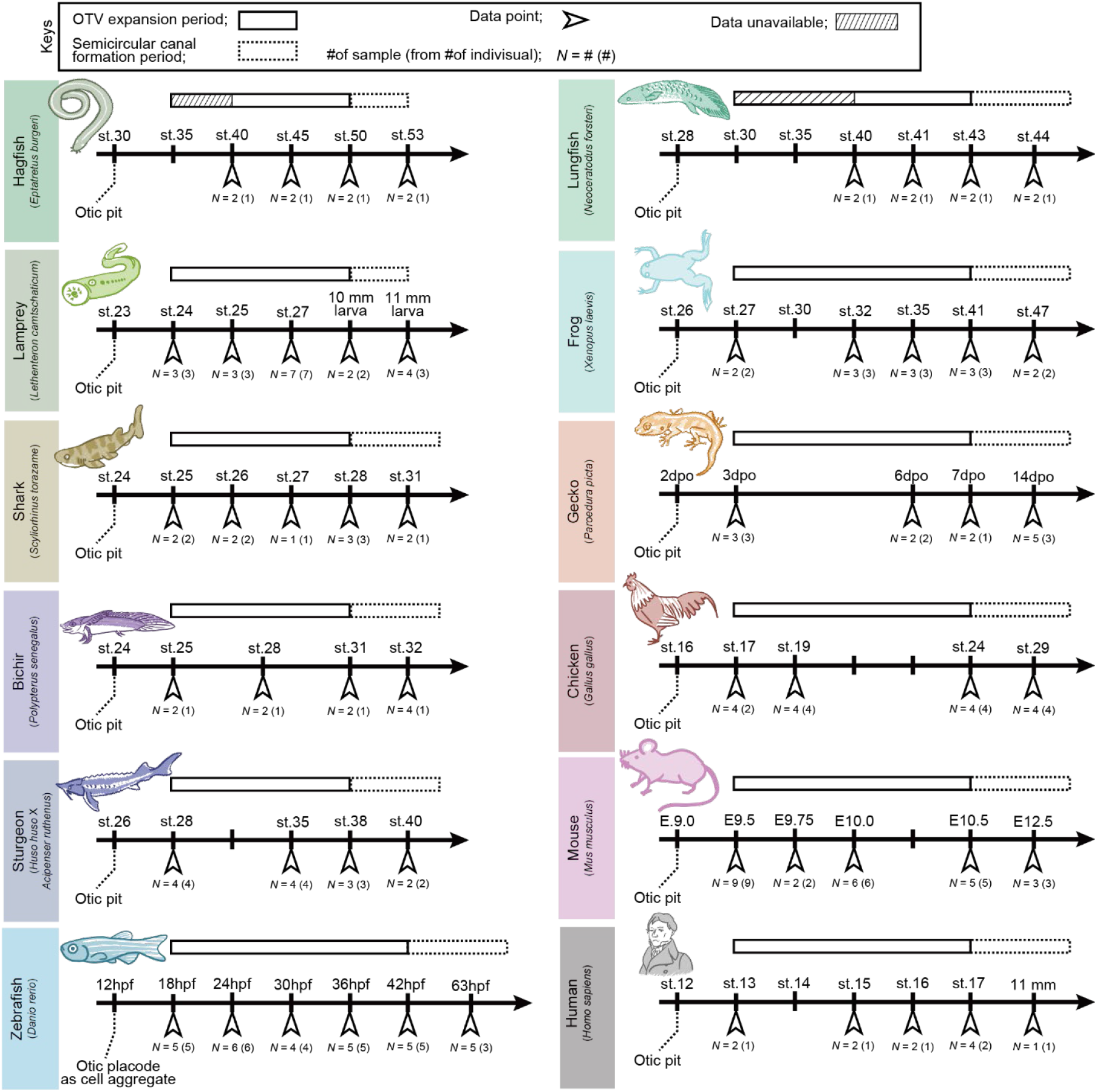
Defining corresponding stages in the inner ear formation in each species. We defined the corresponding periods of inner ear morphogenesis across species. Sampling points and sample sizes used in this study are indicated. The earliest-stage samples of hagfish and lungfish OTVs were partially incomplete due to limited sample accessibility (shaded by diagonal lines).

**Fig. S2.**
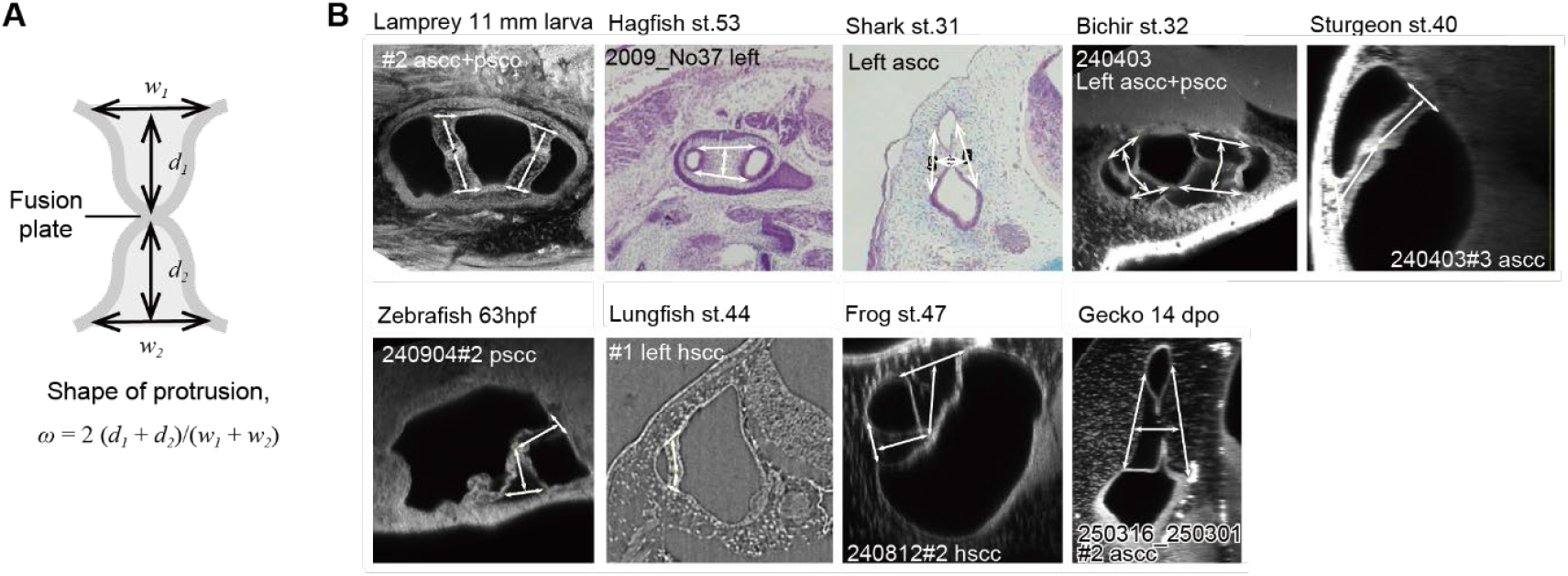
Quantification of the shape of epithelial protrusions across species. **(A)** Definition to quantify the shape of epithelial protrusions in the semicircular canal formation. **(B)** Representative confocal or histological sections containing two protrusions contacting at their tips to form the fusion plate. Overlaid bidirectional arrows indicate the measured regions. Embryos of lampreys, bichirs, sturgeons, zebrafish, frogs, and geckos were stained by RBITC (inverted). Histological sections of hagfish and shark were stained by hematoxylin and eosin. Lungfish embryos were imaged by high-resolution X-ray tomography. Data not shown here were obtained from previously published images (also see table S1). ascc, anterior semicircular canal; hscc, horizontal semicircular canal; pscc, posterior semicircular canal.

**Fig. S3.**
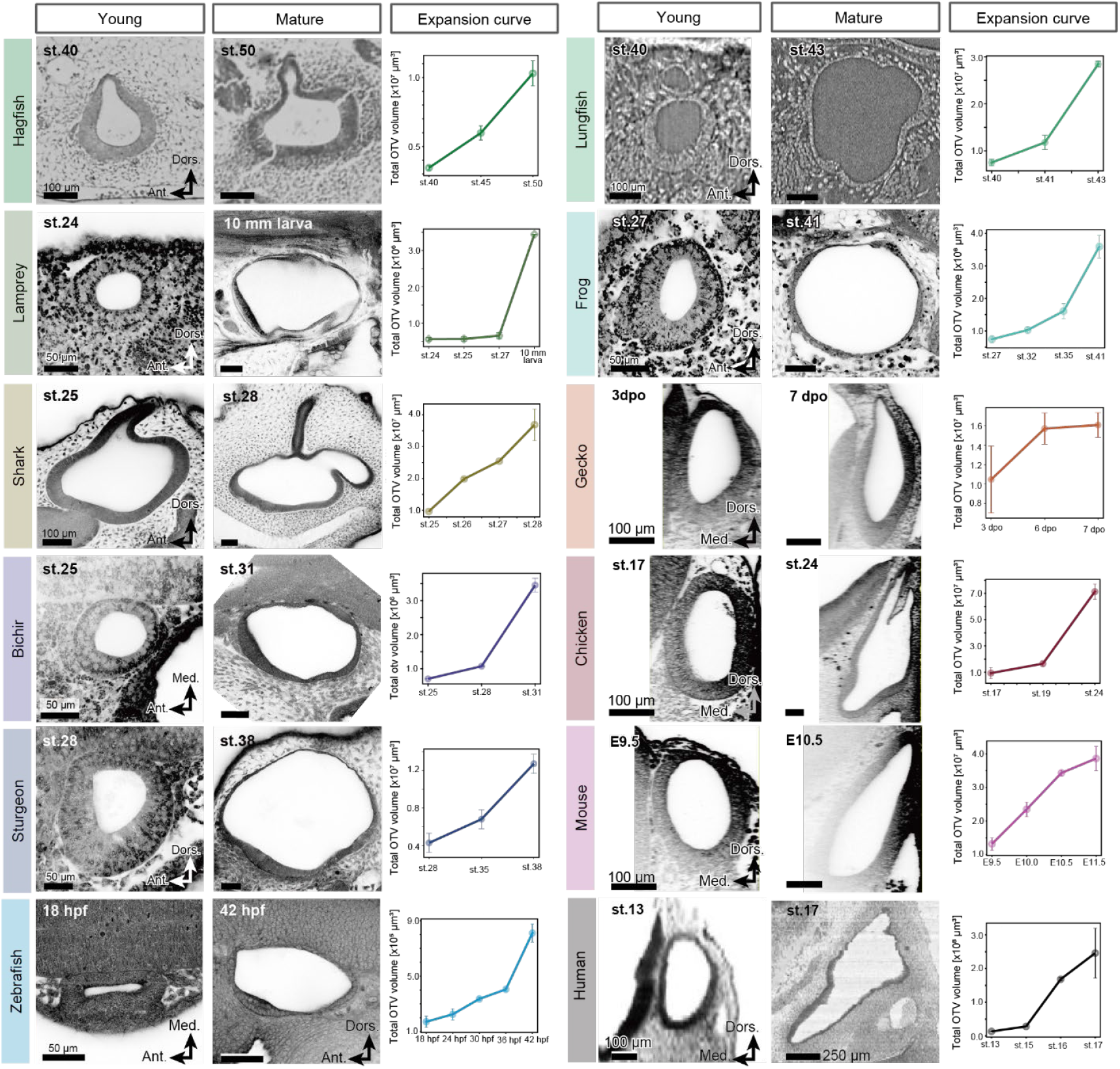
Comparing OTV expansion process across vertebrates. Confocal or histological sectional images of young (left column) and mature (middle column) OTVs are shown in each animal. Lungfish embryos were imaged by high-resolution X-ray tomography. Graphs in right columns are the total OTV size as a function of the developmental stage, showing the OTVs monotonously expand as development progresses in all animals examined in this study. Mean ± SD. Scale bars = 100 μm (hagfish, shark, lungfish, gecko, chicken, mouse, human), 50 μm (lamprey, bichir, sturgeon, zebrafish, frog).

**Fig. S4.**
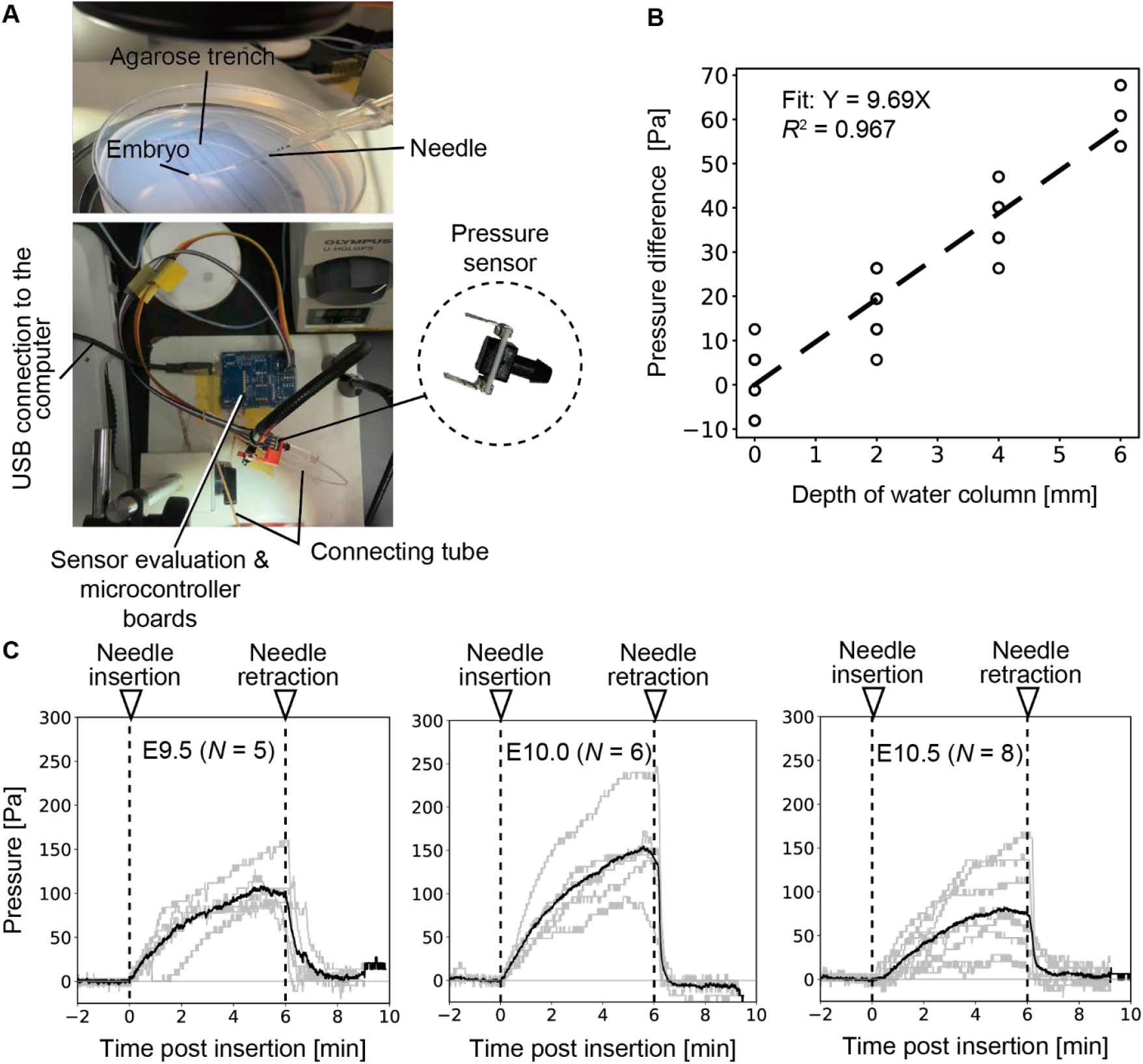
Measurement of luminal pressure in mouse OTV. (**A**) Method and device of pressure measurement. (**B**) Calibration of the pressure sensor using a water column. The dashed line indicates the linear regression fit with coefficient of determination (*R*^*2*^). (**C**) Measurement of luminal pressure in mouse OTV at each developmental stage. Gray lines represent individual measurements and the black line represents the mean value.

**Fig. S5.**
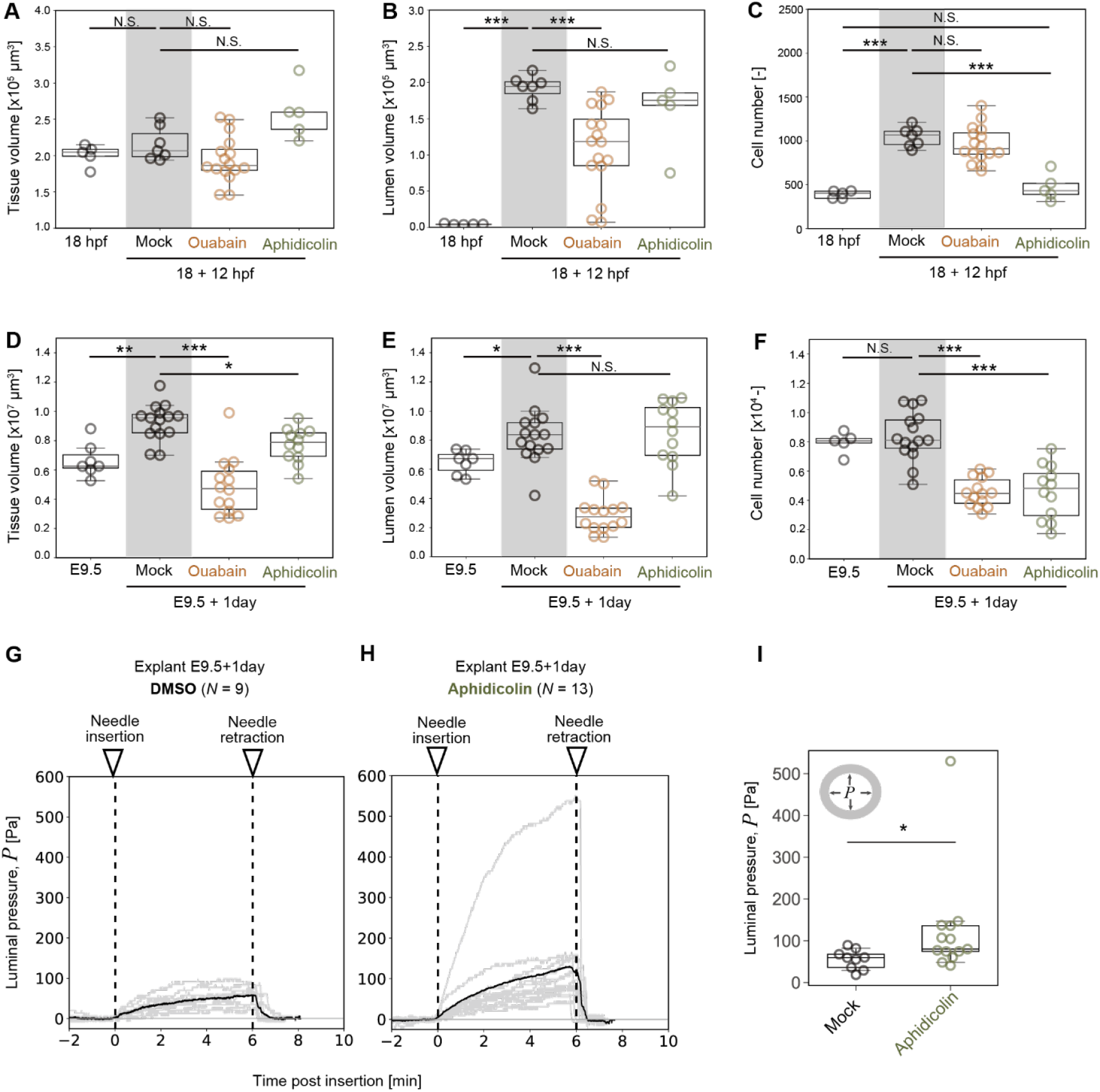
Volume and pressure changes after drug treatments. (**A–C**) Comparisons of volume and cell number among 18 hpf, mock, ouabain, and aphidicolin treated zebrafish embryos. Tissue volume (A), lumen volume (B), and cell number (C) are shown. (**D–F**) Comparisons of volume and cell number among E9.5 mouse embryos, mock, ouabain, and aphidicolin treated explants. Tissue volume (D), lumen volume (E), and cell number (F) are shown. Statistics in (A–F); Games-Howell test. \**P* < 0.05, \*\**P* < 0.01, \*\*\**P* < 0.001, N.S., not significant. (**G, H**) Measurement of the luminal pressure in mouse OTV explants after mock (G) and aphidicolin (H) treatment. Grey and black lines represent individual and mean measurement, respectively. (**I**) Comparison of the luminal pressure in mouse OTV explants after mock and aphidicolin treatment. Brunner-Munzel test. \*\**P*<0.01.

**Fig. S6.**
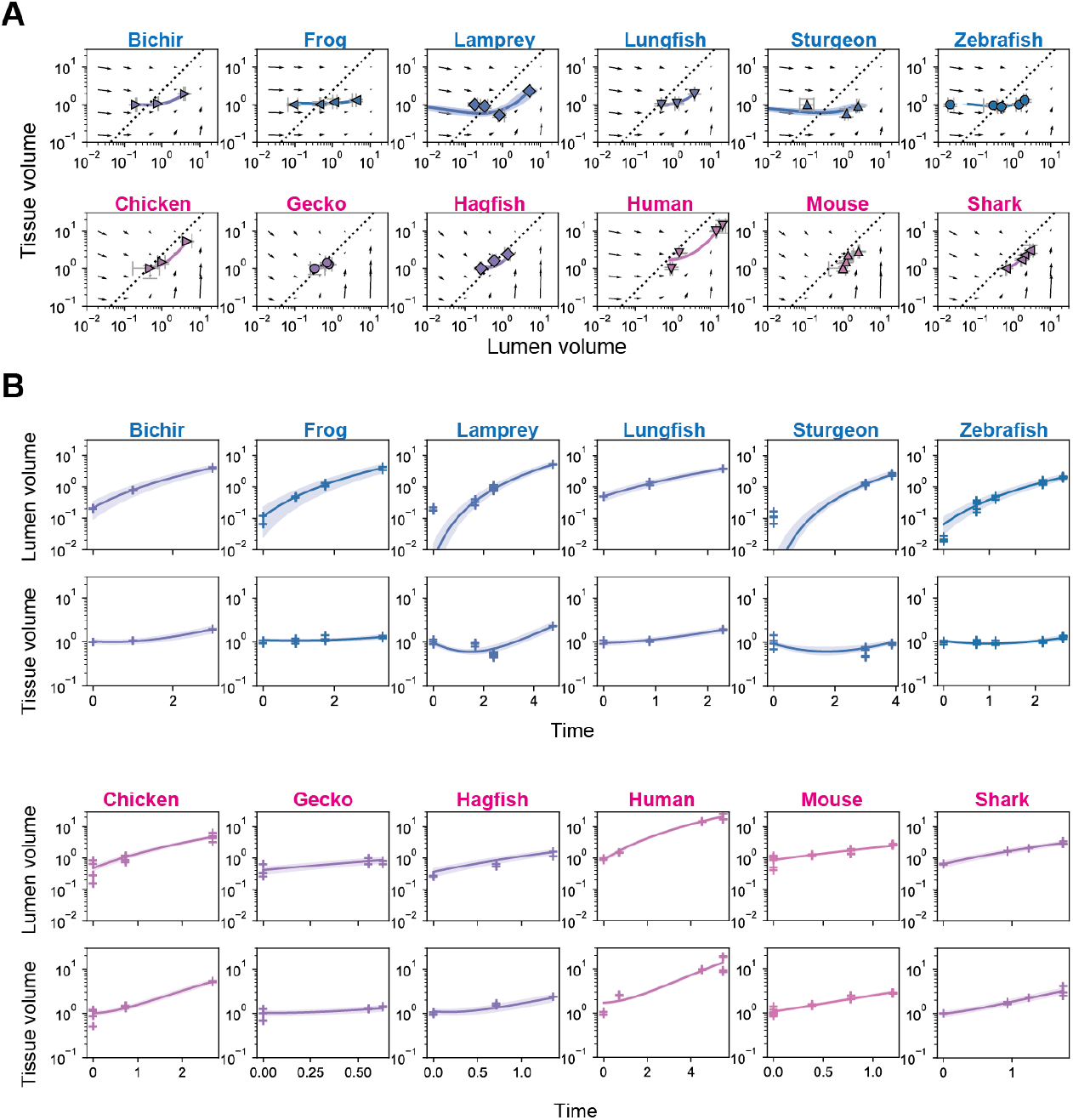
Fit of the mathematical model to the observed data across species. Predicted and observed temporal changes in lumen volume *V*_*L*_ and tissue volume *V*_*E*_ for each species. Volumes are nondimensionalized by the initial tissue volume. (**A**) Temporal trajectories of OTV geometry plotted in the space of lumen and tissue volumes. (**B**) Temporal changes in lumen volume (upper rows) and tissue volumes (lower rows), plotted against time inferred from lumen volume dynamics by setting γ = 1. In both (A) and (B), solid lines and shaded regions indicate the posterior predictive mean and 95% confidence intervals of the mathematical model. In (A), the dotted line denotes the target isometric-scaling relationship approached in the long-time limit of the mathematical model, and small black arrows indicate the vector field of the model evaluated at the posterior mean of the parameters. Points represent the observed data used for parameter estimation. In (B), the observation times are given by the posterior mean estimates of the corresponding latent times.

**Fig. S7.**
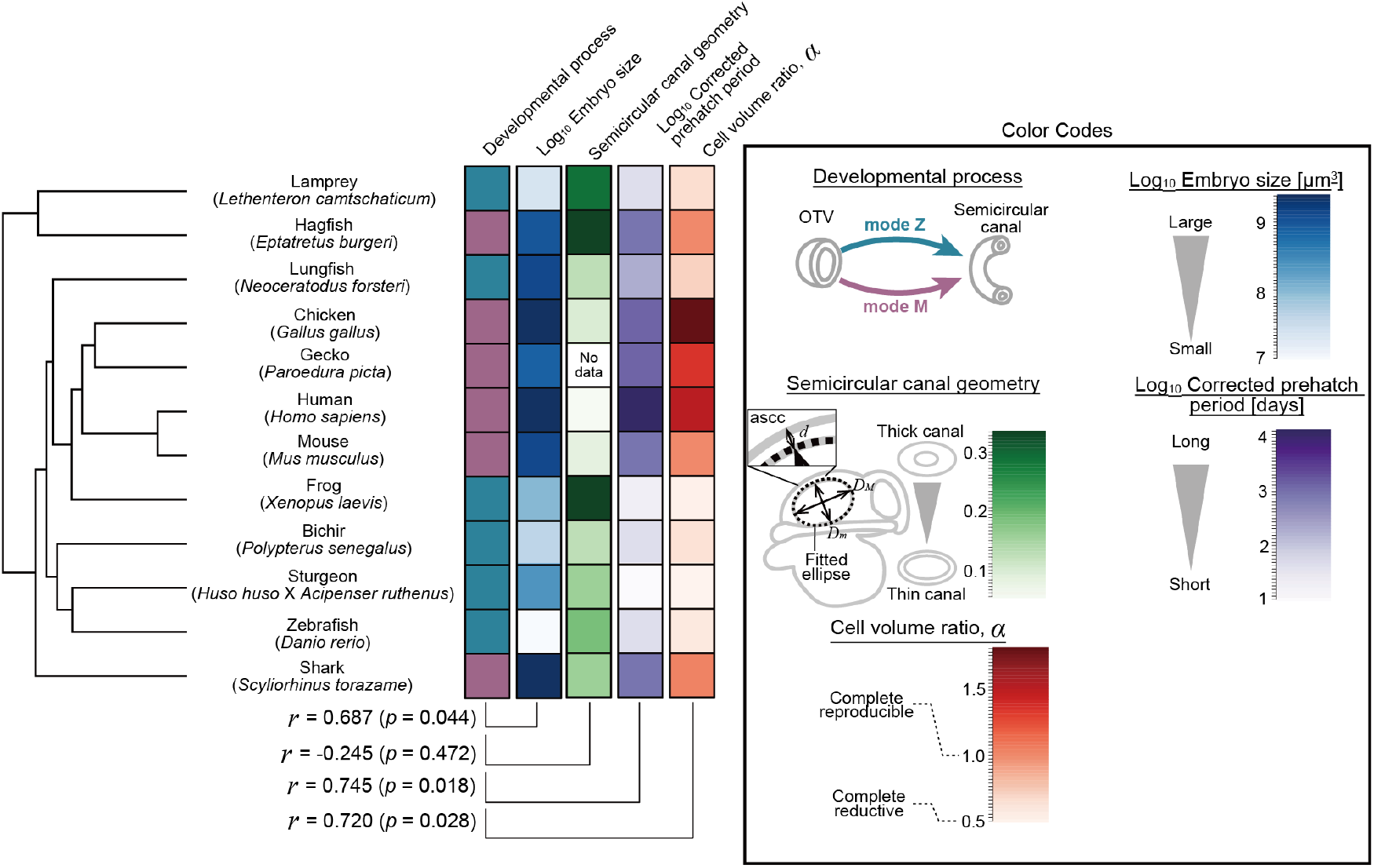
Correlation analysis between the inner ear morphogenetic processes and selected traits. Embryo and adult traits mapped onto each vertebrate species. Each column in the matrix represents individual trait, with color codes shown on the right side. Phylogenetic correlation analysis between the mode of morphogenetic process (discrete trait) and continuous traits was performed. The mean correlation coefficient (*r*) and two-sided posterior tail probability (*p*) were obtained from MCMC sampling. *d*, diameter of the anterior semicircular canal; *D*_*M*_ and *D*_*m*_, diameters of major and minor axis of the fitted ellipse, respectively.

**Fig. S8.**
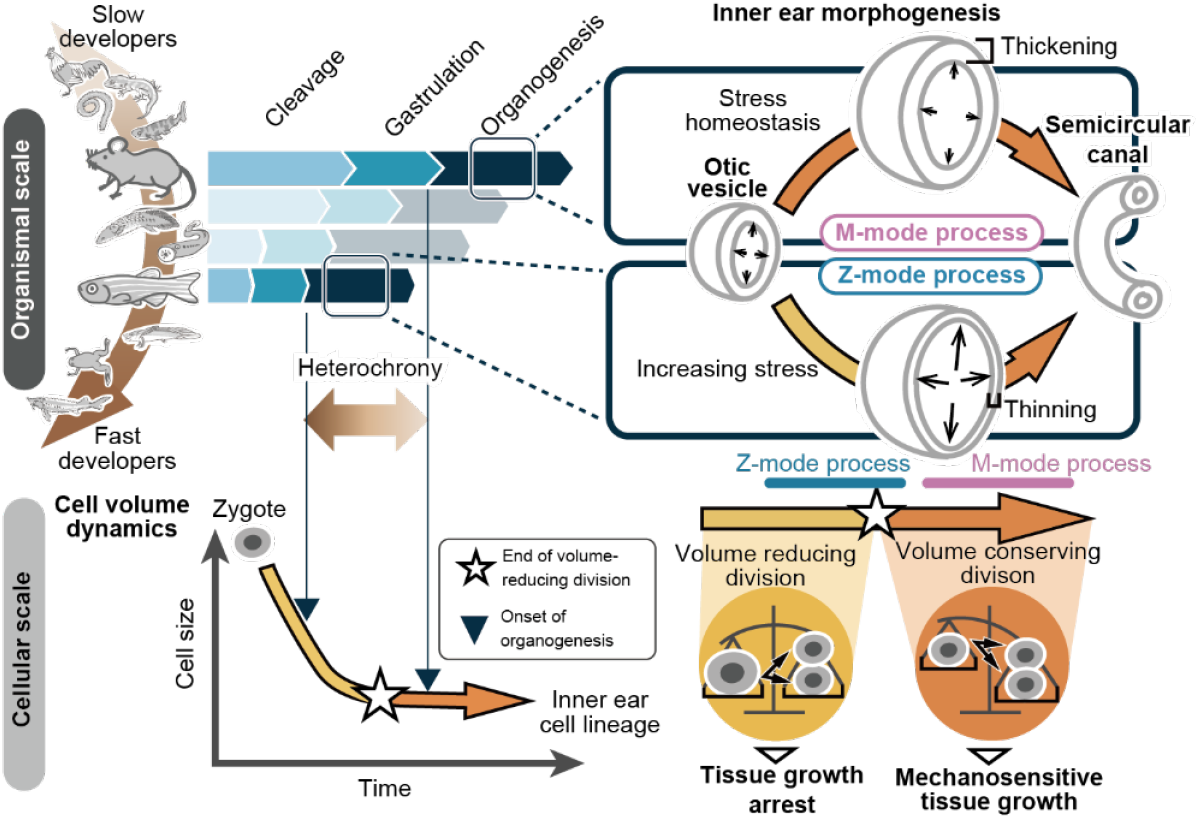
Proposed causal link between evolving developmental duration to divergent inner ear morphogenesis. Evolution of developmental duration is accompanied by heterochrony between the transition in cell-volume dynamics (star) and organogenesis onset (triangles). This heterochrony differentiates the capacity for mechanosensitive tissue growth during organogenesis: fast developers are restricted, whereas slow developers are permitted coupled proliferation and growth. The resulting divergence in tissue growth accessibility during morphogenesis yields distinct OTV expansion mechanics, yet converges on conserved semicircular canal structure.

**Table S1.**
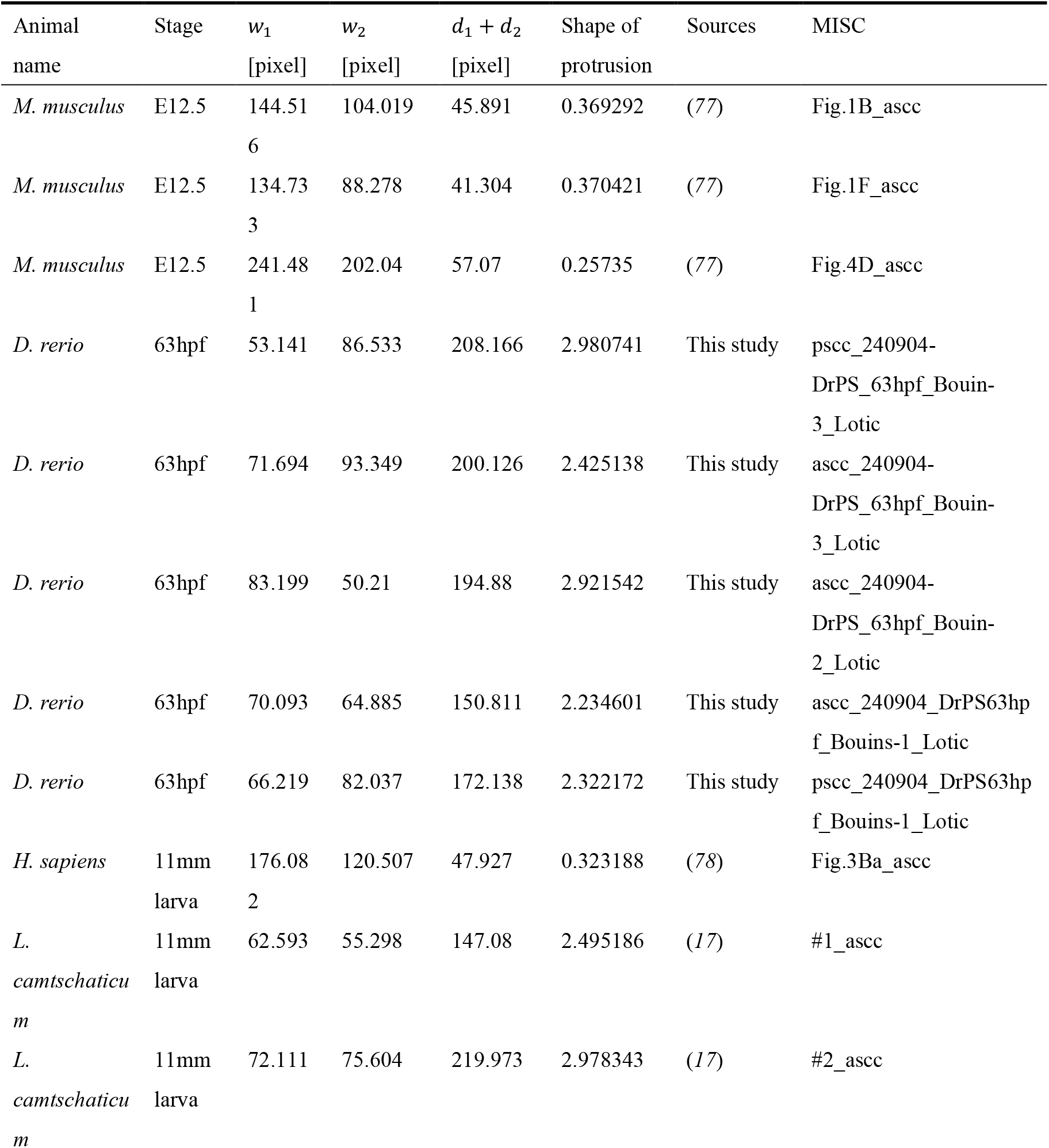
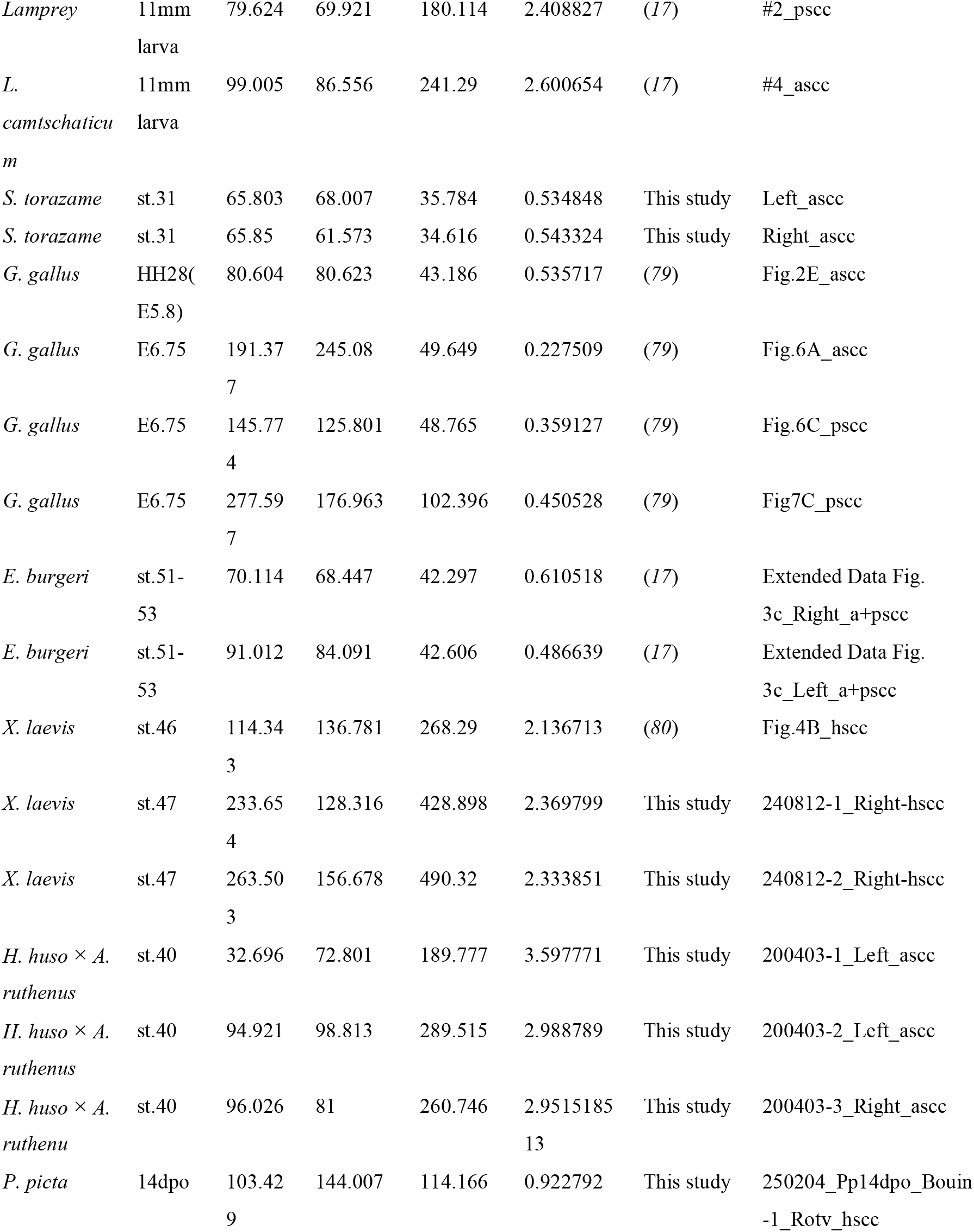
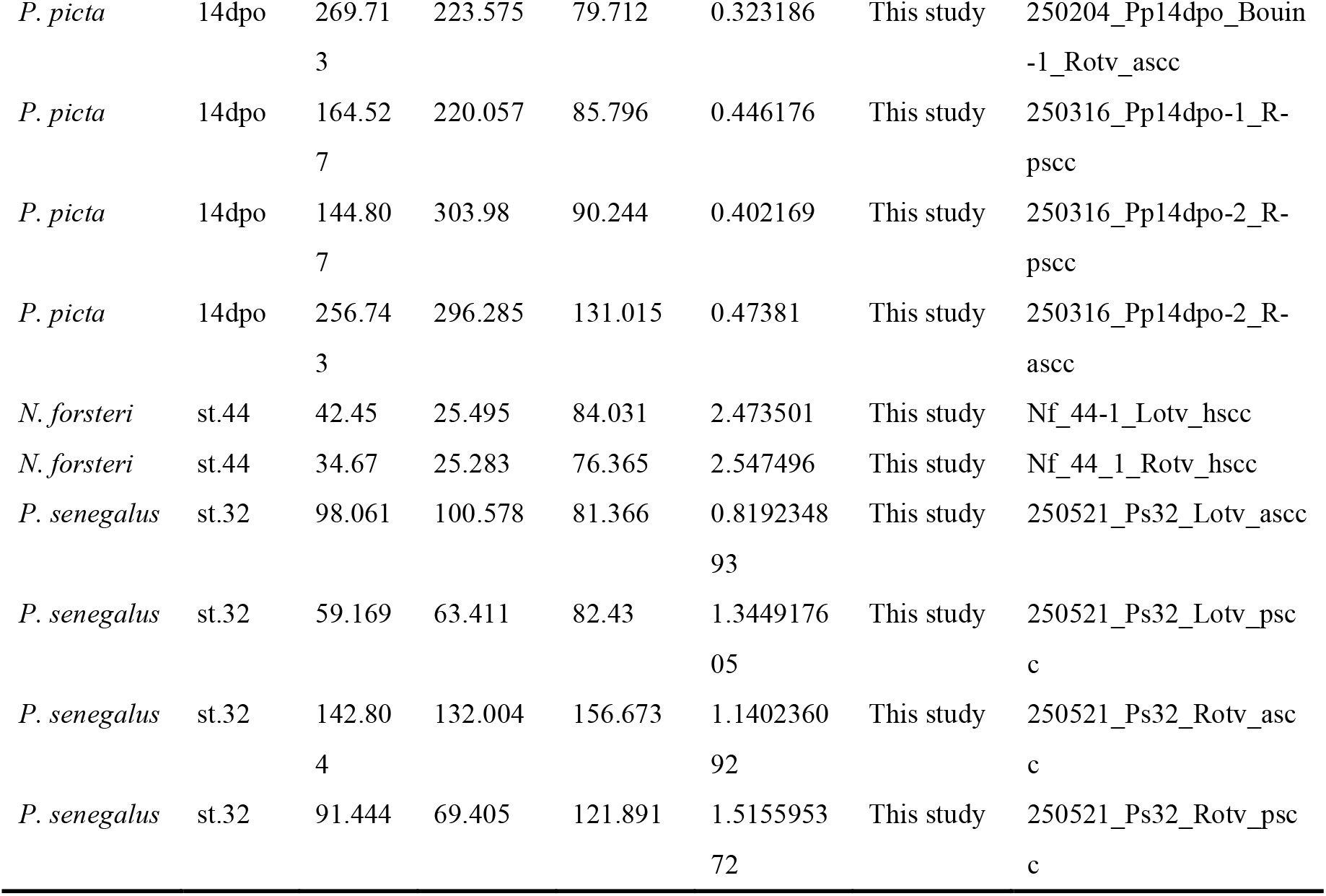
List of measurements of shape of protrusion in the developing inner ear. ascc, anterior semicircular canal; hscc, horizontal semicircular canal; pscc, posterior semicircular canal.

**Table S2.** List of measurements of head volume in each animal.

| Animal name | Sample IDs | Volume [$\mu\text{m}^3$] |
| --- | --- | --- |
| *D. rerio* | 240810DrPS_18hpf_Bouins_RhB_whole-1_head | 14,280,300 |
| *D. rerio* | 240810DrPS_18hpf_Bouins_RhB_whole-2_head | 10,395,200 |
| *D. rerio* | 240627_DrWT18hpf_Bouin-1_head | 12,885,700 |
| *X. laevis* | 240920_Xl_st27_Bouin-3_Rotic_BABB_head | 98,804,300 |
| *X. laevis* | 240920_Xl_st27_Bouin-2_Rotic_BABB_head | 97,212,500 |
| *M. musculus* | 240705_MmE95_Bouin_RhB_head-1_5x | 1,212,770,000 |
| *L. camtschaticum* | 240624_Lcst24_Bouins_RhB_CUBIC_head-1_10x | 32,579,100 |
| *L. camtschaticum* | 240624_Lcst24_Bouins_RhB_CUBIC_head-2_10x | 32,913,000 |
| *L. camtschaticum* | 240624_Lcst24_Bouins_RhB_CUBIC_head-3_10x | 30,487,800 |
| *S. torazame* | 240607_wb25AORI_Bouin-1_head | 2,285,050,000 |
| *H. sapiens* | CS13_CC836 | 1,787,650,000 |
| *E. burgeri* | Eb_no35_st35 | 848,336,000 |
| *G. gallus* | 140424_HH16-17_Serra_4xhead | 1,272,950,000 |
| *G. gallus* | 140424_HH17_Serra_4xhead | 2,440,920,000 |
| *H. huso* × *A. ruthenus* | 250407_bester_st28-3_head_BABB | 253,092,000 |
| *H. huso* × *A. ruthenus* | 250407_bester_st28-4_head_BABB | 260,511,000 |
| *P. picta* | 250208_Pp3dpo_1_Bouin_RhB_head | 565,023,000 |
| *P. picta* | 250208_Pp3dpo_2_Bouin_RhB_head | 602,604,000 |
| *N. forsteri* | Nf_st40_head | 1,009,850,000 |
| *P. senegalus* | 250524_Ps25_250517_bouin_babb_10x_head | 51,404,532 |

**Table S3.** List of measurements of the anterior semicircular canal geometry. ascc, anterior semicircular canal; d, diameter of ascc; *D*_*M*_, length of the major axis of the fitted ellipse; *D*_*m*_, length of the minor axis of the fitted ellipse.

| Animal name | $(D_M + D_m)/2$ [in pixel] | d [in pixel] | ascc relative thickness | Sources |
| --- | --- | --- | --- | --- |
| *D. rerio* | 148.69 | 28.178 | 0.189508373 | (81), Fig.4.2K |
| *X. laevis* | 87.8205 | 28.284 | 0.322066032 | (82), Fig.3L (st.52 tadpole) |
| *M. musculus* | 221.864 | 21.19 | 0.09550896 | (83), Fig.1 |
| *L. camtschaticum* | 155.2905 | 43.382 | 0.279360296 | (84), Fig.4A (represented by *Petromyzon marinus*) |
| *S. torazame* | 112.4745 | 18.601 | 0.165379708 | (85), Fig.1A (represented by *Scyliorhinus canicula*) |
| *H. sapiens* | 215.9835 | 11.662 | 0.053994865 | (86), Fig.1b |
| *E. burgeri* | 202.8425 | 68 | 0.335235466 | (17), Fig.1c |
| *G. gallus* | 142.9385 | 13.928 | 0.097440508 | (87), Fig.4 (E17 embryo) |
| *H. huso* × *A. ruthenus* | 111.64 | 18.385 | 0.164681118 | (85), Fig.1E (represented by *Acipensor sturio*) |
| *N. forsteri* | 120.617 | 15.232 | 0.126284023 | (85), Fig.1C |
| *P. senegalus* | 83.388 | 10.966 | 0.131505732 | (85), Fig.1D (represented by *Polypterus bichir*) |

**Table S4.**
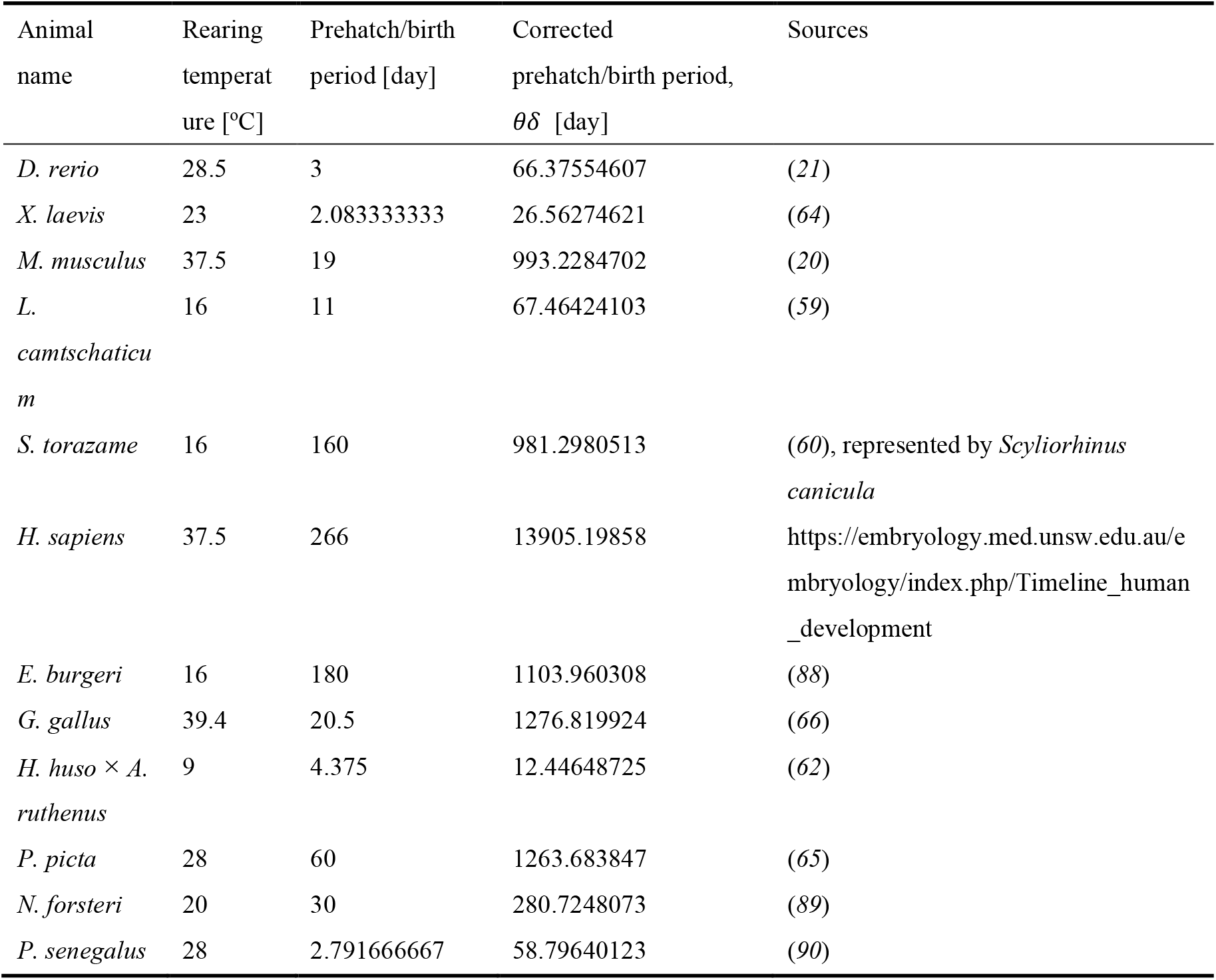
Prehatch developmental period and rearing temperatures collected from staging tables.

| Animal name | Rearing temperature [°C] | Prehatch/birth period [day] | Corrected prehatch/birth period, $\theta\delta$ [day] | Sources |
| --- | --- | --- | --- | --- |
| *D. rerio* | 28.5 | 3 | 66.37554607 | (21) |
| *X. laevis* | 23 | 2.083333333 | 26.56274621 | (64) |
| *M. musculus* | 37.5 | 19 | 993.2284702 | (20) |
| *L. camtschaticum* | 16 | 11 | 67.46424103 | (59) |
| *S. torazame* | 16 | 160 | 981.2980513 | (60), represented by *Scyliorhinus canicula* |
| *H. sapiens* | 37.5 | 266 | 13905.19858 | https://embryology.med.unsw.edu.au/embryology/index.php/Timeline_human_development |
| *E. burgeri* | 16 | 180 | 1103.960308 | (88) |
| *G. gallus* | 39.4 | 20.5 | 1276.819924 | (66) |
| *H. huso* × *A. ruthenus* | 9 | 4.375 | 12.44648725 | (62) |
| *P. picta* | 28 | 60 | 1263.683847 | (65) |
| *N. forsteri* | 20 | 30 | 280.7248073 | (89) |
| *P. senegalus* | 28 | 2.791666667 | 58.79640123 | (90) |

**Movie S1. (separate file)**

Time-lapse movie showing an expanding mouse OTV explant recorded from t=0 to 24 hour.

**Data S1. (separate file)**

List of OTV measurements by developmental stage and species.

**Data S2. (separate file)**

List of pressure measurements in mouse embryos and OTV explants.

**Data S3. (separate file)**

List of OTV measurements after drug treatments in zebrafish and mice.

**Data S4. (separate file)**

List of average cell volume measured along the cell lineage leading to inner ear in zebrafish and mice.

